# epiRomics: a multi-omics R package to identify and visualize enhancers

**DOI:** 10.1101/2021.08.19.456732

**Authors:** Alex M. Mawla, Mark O. Huising

## Abstract

**Summary:** epiRomics is an R package designed to integrate multi-omics data in order to identify and visualize enhancer regions alongside gene expression and other epigenomic modifications. Regulatory network analysis can be done using combinatory approaches to infer regions of significance such as enhancers, when combining ChIP and histone data. Downstream analysis can identify co-occurrence of these regions of interest with other user-supplied data, such as chromatin availability or gene expression. Finally, this package allows for results to be visualized at high resolution in a stand-alone browser.

**Availability and Implementation:** epiRomics is released under Artistic-2.0 License. The source code and documents are freely available through Github (https://github.com/Huising-Lab/epiRomics).

**Supplementary information:** Supplementary data, and methods are available online on biorxiv or *Github*.

## Introduction

The evaluation of the transcriptional landscape between cell types grants the scientific community a deeper understanding of cellular identity, and helps paint the underlying mechanisms that drive phenotype and function (Capobianco, 2014). Bulk RNA sequencing has been a gold standard in the field, followed more recently with the advent of single-omics approaches (Chen, et al., 2019; Conesa, et al., 2016; Kolodziejczyk, et al., 2015; Kulkarni, et al., 2019). However, gene expression represents only a single aspect of what is a sophisticated and interlaced network of genetic and epigenomic regulators that drive and determine cell identity, with perturbations leading to dysfunction and, sometimes, disease (Karczewski and Snyder, 2018).

Chromatin remodeling is a dynamic process that represents one of the epigenetic layers of cell fate maintenance and identity (Andrey and Mundlos, 2017; Muller and Leutz, 2001). Approaches such as DNase I hypersensitive site sequencing (DNASE-Seq) (Boyle, et al., 2008) and Assay for transposase-accessible chromatin using sequencing (ATAC-Seq) (Buenrostro, et al., 2013; Buenrostro, et al., 2015), are commonly used to compare chromatin accessibility between cell types and states. Chromatin immunoprecipitation Sequencing (ChIP-Seq) is another approach used to assess the epigenomic regulators driven by specific transcription factors acting as either activators or suppressors on the genic region, or at distal-intergenic regions, associated with enhancer activity (Daugherty, et al., 2017; de la Torre-Ubieta, et al., 2018; Neph, et al., 2012; Pastor, et al., 2014; Starks, et al., 2019). Alternatively, ChIP-Seq is used in conjunction with antibodies that pull down specific histone modifications associated with regions of chromatin, whose presence can be used to infer whether the region is active, poised or repressed (Calo and Wysocka, 2013; Mellor, 2005). The interrogation of transcription factor binding is not enough to infer transcriptional behavior, as whether or not chromatin is accessible between cell types, and whether the region is active, poised, or repressed, demarked by co-occurrences of histone marks, must be considered in order to fully evaluate the biology (Bemer, 2018; Calo and Wysocka, 2013). Lastly, methylation of DNA, quantifiable through whole-genome bisulfite sequencing analysis (BS-Methyl Seq) (Adusumalli, et al., 2015), can help determine whether accessible chromatin that has recruited the correct histone marks is even available for transcription factor binding and recruitment of co-modulators to drive or suppress gene transcription within cell types (Arand, et al., 2012; He, et al., 2011).

While tools exist to compare these different layers in a pairwise manner, a gap exists to integrate multiple omics layers quickly, and easily to generate high resolution visuals in order to derive more biological meaning behind results. We developed a novel ‘epigenomics in R’, epiRomics, package to solve this issue. We designed epiRomics to accept either browser extensible data (BED) (Kent, et al., 2002) or bigwig (Kent, et al., 2010) files as input for any of the aforementioned types of data. Inclusion of functional annotations, *i.e*. FANTOM (Lizio, et al., 2017), single nucleotide polymorphism (SNP) data from GWAS (Wang, et al., 2010; Zheng, et al., 2015), or Ultra Conserved Non Coding Elements (UCNEs) (Dimitrieva and Bucher, 2012) is also possible – in order to more fully integrate many slices of the genetic and epigenetic pie.

## Functions

epiRomics takes in a user-submitted comma-separated values (csv) file containing hard paths to all BED or bigwig formatted files, optional hexadecimal (hex) color code associations for each file and a user-defined label to group each input data set (*e.g*., ChIP, ATAC, RNA, functional, etc.). The *epiRomics_build_dB* command quickly generates a comprehensive and easily accessible variable of the class “epiRomics-class” containing a GenomicRanges (*GRanges)* object (Lawrence, et al., 2013) that tracks each of these submitted data, along with all other data related to the species, pulled automatically from the UCSC genome database (Kent, et al., 2002). The epiRomics-class variable can easily be integrated with other packages, and the user can also save these data in a csv format for further manual exploration in excel, or other comparable third-party tools.

Putative enhancers can efficiently be called, and then categorized separately – active, poised, repressed, etc., through the *epiRomics_putative_enhancers* function, which will consider user provided histone data. For example, the histone marks H3K27ac and H3K4me1 are commonly used to demark active enhancer regions (Calo and Wysocka, 2013; Creyghton, et al., 2010; Spicuglia and Vanhille, 2012; Spitz and Furlong, 2012) outputting an epiRomics-class variable for downstream use within the package, or outside. This variable can be used further to identify key enhanceosome regions with evidence of co-binding of multiple user-selected transcription factors by implementing the *epiRomics_putative_enhanceosome* command. These data can also be filtered against functional data annotations, such as methylation calls, FANTOM, SNP regions, or UCNEs, through the use of *epiRomics_putative_enhancers_filtered*. A side function is provided within the package making use of decision trees (Kingsford and Salzberg, 2008) in order to classify which transcription factors were most meaningfully associated with different enhancer types, through the use of *epiRomics_predictors*.

For visualization of these differently classified regions, and integration with bigwig data such as gene expression or chromatin availability between cell types, the tool *epiRomics_track_layer* can be used. This makes use of the package *GViz (Hahne and Ivanek, 2016)* to generate resolution, publication-quality encapsulated postscript (eps) files. Specific calls for enhancer regions provided by *epiRomics_putative_enhanceosome* can be visualized. Conversely, if users have specific regions or genes of interest they wish to evaluate, they can do so using *epiRomics_region_of_interest*.

These tools were designed to allow biological relevance to be determined from the integrated multi-omics data that is available for a particular tissue or cell type. For example, a common enhancer region may be present between cell types of a common progenitor, with chromatin accessible across all cell types, methylation may block activity in one cell type, but not the other. Drug treatment, or healthy versus diseased comparisons can quickly be made, and the multitude of SNPs amassed via GWAS can be seamlessly connected to narrow in on deleterious variants that may contribute to disease.

## Results

epiRomics is developed as an R package to be made available through Bioconductor (Gentleman, et al., 2004), and is available under Artistic-2.0 License. epiRomics is designed to integrate a multitude of -omics data – in either BED or bigwig format – in order to identify regions of regulatory interest, such as enhancers, and provide sophisticated, high quality resolution visuals in EPS format for use in publications. Users with little programming experience can use epiRomics to encode colors for individual tracks, cross-reference diverse types of -omics data – such as ATAC- and RNA-Seq, and produce strong candidate lists for putative enhancers common or unique to cell types. Finally, epiRomics is easy to use, with a full walkthrough with sample data available through its companion vignette.

## Supporting information

Methods for Aggregated Data in Vignette

Aggregated Datasets Accession List

Getting Started with EpiRomics Vignette

EpiRomics Manual

