## Supplementary material for "epiRomics: a multi-omics R package to identify and visualize enhancers": Methods for Aggregated Data in Vignette

**Sample Data Methods**

***ATAC-sequencing from human beta and alpha cells***

Raw fastq files for all human alpha and beta samples were retrieved from GEO accession GSE76268 (Ackermann, et al., 2016). Samples were processed using the ENCODE-DCC ATAC sequencing pipeline, aligning to the hg38 (Harrow, et al., 2012) build of the human genome (Consortium, 2012; Davis, et al., 2018). Peak calls generated through the pipeline using MACS2 (Zhang, et al., 2008) were analyzed downstream through the BioConductor package DiffBind (Ross-Innes, et al., 2012) in order to identify differentially enriched chromatin regions between the two cell types. Bigwigs merged by cell type generated through the pipeline were subsetted to chromosome 1 using UCSC command line tools (Kent, et al., 2010).

***RNA-sequencing from human beta and alpha cells***

Raw, companion fastq files for all human alpha and beta samples were also retrieved from GEO accession GSE76268 (Ackermann, et al., 2016). Fastq files were quality controlled using the tool fastp (Chen, et al., 2018), and aligned using STAR (Dobin, et al., 2013) to the hg38 build of the human genome. Wiggle files produced by the STAR aligner were then merged by cell type using UCSC command line tools. Bigwigs merged by cell type generated through the pipeline were subsetted to chromosome 1 using UCSC command line tools.

***ChIP- and histone-sequencing data***

ChIP-sequencing peak calls generated using MACS2 for human pancreatic islet transcription factors Foxa2, MafB, Nkx2.2, Nkx6.1, and Pdx1 were retrieved from the EMBL-EBI repository database E-MTAB-1919 (Pasquali, et al., 2014). All peak calls were lifted over to the hg38 genome build using the UCSC genome browser liftOver tool (Kent, et al., 2002).

Histone-sequencing peak calls generated using MACS2 for histones H3k27ac and H3k4me1 were retrieved from GEO accession GSE16256 (Bernstein, et al., 2010), and for histone H2A.Z from the EMBL-EBI repository database E-MTAB-1919 (Pasquali, et al., 2014). All peak calls were lifted over to the hg38 genome build using the UCSC genome browser liftOver tool.

***Incorporated functional annotation***

The FANTOM5 human enhancer database (Lizio, et al., 2015) was retrieved, and all regions were lifted over to the hg38 genome build using the UCSC genome browser liftOver tool.

Human ultra-conserved non-coding elements (UCNEs) were retrieved form the UCNE database (Dimitrieva and Bucher, 2012), and all regions were lifted over to the hg38 genome build using the UCSC genome browser liftOver tool.

The human islet regulome database was retrieved and lifted over to the hg38 genome build using the UCSC genome browser liftOver tool as well (Miguel-Escalada, et al., 2019). The super enhancer functional annotation was defined based on the authors’ calls. The active enhancers annotation was filtered and selected from the authors’ supplemental dataset, in order to remove other calls that were not deemed ‘active enhancers’.
