## Supplementary material for "epiRomics: a multi-omics R package to identify and visualize enhancers": Getting Started with EpiRomics Vignette

Authors: Alex M. Mawla & Mark O. Huising. Copyright 2020 - Present.

2021-10-05

#### Contents

**Abstract** 1

**Citation** 2

**Loading the epiRomics package and dependencies for vignette** 2

**Brief explanation of example data** 2

**How to load and build the database** 3

**Delineating active enhancers using H3k4me1 and H3k27ac marks as a proxy** 4

**Cross-referencing enhancer calls to other databases** 5

**Screening for high transcription factor co-binding sites** 7

**Transcription factor decision trees** 9

**Intersecting and visualizing ATAC- and RNA-Seq data** 10

**Session Information** 29

#### Abstract

**Summary** epiRomics is an R package designed to integrate multi-omics data in order to identify and visualize enhancer regions alongside gene expression and other epigenomic modifications. Regulatory network analysis can be done using combinatory approaches to infer regions of significance such as enhancers, when combining ChIP and histone data. Downstream analysis can identify co-occurrence of these regions of interest with other user-supplied data, such as chromatin availability or gene expression. Finally, this package allows for results to be visualized at high resolution in a stand-alone browser.

**Availability and Implementation** epiRomics is released under Artistic-2.0 License. The source code and documents are freely available through Github (https://github.com/Huising-Lab/epiRomics).

**Contact**

**Supplementary information** Supplementary data, and methods are available online on *biorXiv* or *Github.*

**Competing Interest Statement**

The authors have declared no competing interest.

### Citation

If you use epiRomics in published research, please cite:

Mawla, AM& Huising, MO. **epiRomics: a multi-omics R package to identify and visualize enhancers.**

biorXiv 2021. doi:https://doi.org/10.1101/2021.08.19.456732

### Loading the epiRomics package and dependencies for vignette

```
## loading packages

library(epiRomics)

library(TxDb.Hsapiens.UCSC.hg38.knownGene)

library(org.Hs.eg.db)
```

### Brief explanation of example data

This package includes some example data to get you started, delineating human pancreatic islet enhancers between alpha and beta cells.

Human pancreatic islet alpha and beta ATAC- and companion RNA- Seq data were retrieved from GEO accession GSE76268 (Ackermann, et al., 2016).

ATAC samples were processed using the ENCODE-DCC ATAC sequencing pipeline, aligning to the hg38 (Harrow, et al., 2012) build of the human genome (Consortium, 2012; Davis, et al., 2018).

Peak calls generated through the pipeline using MACS2 (Zhang, et al., 2008) were analyzed downstream through the BioConductor package DiffBind (Ross-Innes, et al., 2012) in order to identify differentially enriched chromatin regions between the two cell types.

RNA samples were quality controlled using the tool fastp (Chen, et al., 2018), and aligned using STAR (Dobin, et al., 2013) to the hg38 build of the human genome. Wiggle files produced by the STAR aligner were then merged by cell type using UCSC command line tools.

Bigwigs merged by cell type were subsetted to chromosome 1 using UCSC command line tools (Kent, et al., 2010).

ChIP-sequencing peak calls generated using MACS2 for human pancreatic islet transcription factors Foxa2, MafB, Nkx2.2, Nkx6.1, and Pdx1 were retrieved from the EMBL-EBI repository database E-MTAB-1919 (Pasquali, et al., 2014). All peak calls were lifted over to the hg38 genome build using the UCSC genome browser liftOver tool (Kent, et al., 2002).

Histone-sequencing peak calls generated using MACS2 for histones H3k27ac and H3k4me1 were retrieved from GEO accession GSE16256 (Bernstein, et al., 2010), and for histone H2A.Z from the EMBL-EBI repository database E-MTAB-1919 (Pasquali, et al., 2014). All peak calls were lifted over to the hg38 genome build using the UCSC genome browser liftOver tool.

The FANTOM5 human enhancer database (Lizio, et al., 2015) was retrieved, and all regions were lifted over to the hg38 genome build using the UCSC genome browser liftOver tool.

Human ultra-conserved non-coding elements (UCNEs) were retrieved form the UCNE database (Dimitrieva and Bucher, 2012), and all regions were lifted over to the hg38 genome build using the UCSC genome browser liftOver tool.

The human islet regulome database was retrieved (Miguel-Escalada, et al., 2019) and all regions were lifted over to the hg38 genome build using the UCSC genome browser liftOver tool.

#### How to load and build the database

Lets load and take a look at how to properly format the datasets epiRomics uses to build the initial database.

```
example_epiRomics_Db_sheet <- read.csv(
  file = system.file(
    "extdata",
    "example_epiRomics_Db_sheet_user_paths.csv",
    package = "epiRomics"
  )
)

## Required columns are: name, path, genome, format, and type

## The genome must also be in proper format, e.g. mm10 or hg38

## Type of data can be histone, methyl, SNP, or ChIP.
## ChIP is required for some downstream functions to work appropriately.

## Not run
#head(example_epiRomics_Db_sheet)
```

epiRomics_build_dB constructs a database of class epiRomics with this data sheet

```
epiRomics_dB <- epiRomics_build_dB(
  epiRomics_db_file =
    system.file(
      "extdata",
      "example_epiRomics_Db_sheet_user_paths.csv",
      package = "epiRomics"
    ),
  txdb_organism =
```

```
    "TxDb.Hsapiens.UCSC.hg38.knownGene::TxDb.Hsapiens.UCSC.hg38.knownGene",
  epiRomics_genome = "hg38",
  epiRomics_organism = "org.Hs.eg.db"
)
#> Building enhancers...
#> snapshotDate(): 2021-05-18
#> loading from cache
#> 'select()' returned 1:1 mapping between keys and columns
#> Building promoters...
#> Building 1to5kb upstream of TSS...
#> Building intergenic...
#> Building cds...
#> Building 5UTRs...
#> Building 3UTRs...
#> Building exons...
#> Building first exons...
#> Building introns...
#> Building intron exon boundaries...
#> Building exon intron boundaries...
#> Building CpG islands...
#> Building CpG shores...
#> Building CpG shelves...
#> Building inter-CpG-islands...
#> snapshotDate(): 2021-05-18
#> Building lncRNA transcripts...
#> loading from cache
```

#### Delineating active enhancers using H3k4me1 and H3k27ac marks as a proxy

There is a lot of flexibility for data exploration here. In this example, we search for putative enhancers using two histone marks known to co-occur at enhancer regions - h3k4me1 & h3k27ac

```
epiRomics_putative_enhancers <-
  epiRomics_enhancers(
    epiRomics_dB,
    epiRomics_histone_mark_1 =
      "h3k4me1",
    epiRomics_histone_mark_2 = "h3k27ac"
  )

## Taking a look, we see a  list of 19,692 putative enhancers

epiRomics_putative_enhancers@annotations
#> GRanges object with 19692 ranges and 0 metadata columns:
#>           seqnames            ranges strand
#>              <Rle>         <IRanges>  <Rle>
#>       [1]     chr1    999886-1000011      *
#>       [2]     chr1   1000228-1000811      *
#>       [3]     chr1   1000850-1001468      *
#>       [4]     chr1   1005007-1006023      *
```

```
#>        [5]      chr1   1013701-1013893      *
#>        ...       ...               ...    ...
#>    [19688]      chrY 12392544-12392994      *
#>    [19689]      chrY 13282680-13282760      *
#>    [19690]      chrY 15455449-15455788      *
#>    [19691]      chrY 19066496-19066508      *
#>    [19692]      chrY 19075542-19075899      *
#>   -------
#>   seqinfo: 595 sequences (1 circular) from hg38 genome
```

### Cross-referencing enhancer calls to other databases

#### FANTOM Enhancer Database

Now we have a list of regions as possible candidates for enhancers, but where do we go from here? One way to increase confidence of these calls is to cross this list against an enhancer database, for instance, FANTOM.

```
## NOTE: This option may not be available for all organisms.

epiRomics_putative_enhancers_filtered_fantom <-
  epiRomics_enhancers_filter(epiRomics_putative_enhancers, epiRomics_dB,
                             epiRomics_type =
                               "hg38_custom_fantom")

## Taking a look, we see a reduced number of 2,749 candidate regions

epiRomics_putative_enhancers_filtered_fantom@annotations
#> GRanges object with 2749 ranges and 0 metadata columns:
#>          seqnames              ranges strand
#>             <Rle>           <IRanges>  <Rle>
#>      [1]     chr1     1021242-1021277      *
#>      [2]     chr1     1021318-1021698      *
#>      [3]     chr1     1079632-1080061      *
#>      [4]     chr1     1080101-1080628      *
#>      [5]     chr1     1128200-1128445      *
#>      ...      ...                 ...    ...
#>   [2745]     chrX 154369950-154370183      *
#>   [2746]     chrX 154371971-154372237      *
#>   [2747]     chrX 154372350-154372695      *
#>   [2748]     chrX 154517139-154517596      *
#>   [2749]     chrX 154734550-154734738      *
#>   -------
#>   seqinfo: 595 sequences (1 circular) from hg38 genome
```

#### Human Pancreatic Islet Regulome Enhancer Database

We can also filter putative enhancer calls against active enhancers from the human islet regulome database

```
epiRomics_putative_enhancers_filtered_regulome_active <-
  epiRomics_enhancers_filter(epiRomics_putative_enhancers,
```

```
                             epiRomics_dB,
                             epiRomics_type = "hg38_custom_regulome_active")

epiRomics_putative_enhancers_filtered_regulome_active@annotations
#> GRanges object with 6025 ranges and 0 metadata columns:
#>          seqnames              ranges strand
#>             <Rle>           <IRanges>  <Rle>
#>      [1]     chr1     1068896-1068951      *
#>      [2]     chr1     1069171-1069333      *
#>      [3]     chr1     1079632-1080061      *
#>      [4]     chr1     1080101-1080628      *
#>      [5]     chr1     1158358-1158930      *
#>      ...      ...                 ...    ...
#>   [6021]     chrX 153381411-153381523      *
#>   [6022]     chrX 153381677-153381956      *
#>   [6023]     chrX 153382322-153382448      *
#>   [6024]     chrX 153985442-153985689      *
#>   [6025]     chrX 154091801-154091996      *
#>   -------
#>   seqinfo: 595 sequences (1 circular) from hg38 genome
```

##### Human Pancreatic Islet Regulome Super-Enhancer Database

We can also filter putative enhancer calls against super enhancers from human islet regulome database

```
epiRomics_putative_enhancers_filtered_regulome_super <-
  epiRomics_enhancers_filter(epiRomics_putative_enhancers,
                             epiRomics_dB, epiRomics_type = "hg38_custom_regulome_super")

epiRomics_putative_enhancers_filtered_regulome_super@annotations
#> GRanges object with 2401 ranges and 0 metadata columns:
#>          seqnames            ranges strand
#>             <Rle>         <IRanges>  <Rle>
#>      [1]     chr1   7574092-7574479      *
#>      [2]     chr1   7574640-7575094      *
#>      [3]     chr1   8169274-8169689      *
#>      [4]     chr1   8170112-8170857      *
#>      [5]     chr1   8174089-8174358      *
#>      ...      ...               ...    ...
#>   [2397]    chr22 46109916-46110442      *
#>   [2398]    chr22 46115774-46116154      *
#>   [2399]    chr22 46116326-46116501      *
#>   [2400]     chrX 39813348-39813627      *
#>   [2401]     chrX 39814304-39814607      *
#>   -------
#>   seqinfo: 595 sequences (1 circular) from hg38 genome
```

##### Human Ultra-Conserved Non-Coding Elements Database

We can also filter putative enhancer calls against Ultra-Conserved Non-Coding Elements

```r
epiRomics_putative_enhancers_filtered_ucnes <-
  epiRomics_enhancers_filter(epiRomics_putative_enhancers,
                             epiRomics_dB,
                             epiRomics_type = "hg38_custom_ucnes")

epiRomics_putative_enhancers_filtered_ucnes@annotations
#> GRanges object with 11 ranges and 0 metadata columns:
#>        seqnames              ranges strand
#>           <Rle>           <IRanges>  <Rle>
#>    [1]     chr1 164635220-164635921      *
#>    [2]     chr1 164711914-164712296      *
#>    [3]     chr1 164712350-164713071      *
#>    [4]     chr1 200079185-200079426      *
#>    [5]     chr1 213585694-213586385      *
#>    [6]     chr3   71131859-71132164      *
#>    [7]     chr9 106921420-106921764      *
#>    [8]    chr11 114163425-114164860      *
#>    [9]    chr15   36903894-36904085      *
#>   [10]    chr15   53447393-53447809      *
#>   [11]    chr21   16534340-16534665      *
#>   -------
#>   seqinfo: 595 sequences (1 circular) from hg38 genome
```

#### Screening for high transcription factor co-binding sites

Biology has established that enhancers can be quite redundant, and not all play an active role in regulating a cell's activity. How can we utilize other epigenomic data in order to identify true enhanceosome regions? One way is to cross this list against all ChIP data of the cell type. A true enhanceosome region should have made it through our filtering thus far, and contain several binding sites for known TFs. Co-binding is expected, and the list is sorted by the highest number of ChIP hits within the region.

```r
epiRomics_putative_enhanceosome_fantom <-
  epiRomics_enhanceosome(epiRomics_putative_enhancers_filtered_fantom,
                         epiRomics_dB)
#> >> preparing features information...      2021-10-05 17:43:33
#> >> identifying nearest features...        2021-10-05 17:43:34
#> >> calculating distance from peak to TSS...   2021-10-05 17:43:35
#> >> assigning genomic annotation...        2021-10-05 17:43:35
#> >> adding gene annotation...          2021-10-05 17:44:15
#> 'select()' returned 1:many mapping between keys and columns
#> >> assigning chromosome lengths           2021-10-05 17:44:15
#> >> done...                    2021-10-05 17:44:15

## Taking a look, we see the top candidates meet the criteria we list as expected

epiRomics_putative_enhanceosome_fantom@annotations
#> GRanges object with 2749 ranges and 19 metadata columns:
#>        seqnames              ranges strand |     foxa2      mafb    nkx2_2
#>           <Rle>           <IRanges>  <Rle> | <integer> <integer> <integer>
#>    183     chr1 154418514-154419684      * |         2         2         1
#>   1096     chr9     2242369-2242873      * |         2         1         1
```

```
#>   2615     chr22   30310745-30311570      * |         2         1         2
#>     34      chr1   10685395-10688670      * |         1         0         1
#>    792      chr6   30748438-30749427      * |         2         1         1
#>    ...       ...                 ...    ... .       ...       ...       ...
#>   2743      chrX 153927339-153927701      * |         0         0         0
#>   2745      chrX 154369950-154370183      * |         0         0         0
#>   2746      chrX 154371971-154372237      * |         0         0         0
#>   2747      chrX 154372350-154372695      * |         0         0         0
#>   2748      chrX 154517139-154517596      * |         0         0         0
#>             nxk6_1      pdx1      h2az ChIP_Hits             annotation   geneChr
#>          <integer> <integer> <integer> <numeric>            <character> <integer>
#>    183           2         1         2        10 Intron (ENST00000622..         1
#>   1096           1         2         1         8      Distal Intergenic         9
#>   2615           1         1         1         8      Promoter (2-3kb)        22
#>     34           1         2         2         7 Intron (ENST00000377..         1
#>    792           1         1         1         7 Intron (ENST00000656..         6
#>    ...         ...       ...       ...       ...                    ...       ...
#>   2743           0         0         0         0      Promoter (<=1kb)        23
#>   2745           0         0         0         0      Promoter (1-2kb)        23
#>   2746           0         0         0         0      Promoter (<=1kb)        23
#>   2747           0         0         0         0      Promoter (1-2kb)        23
#>   2748           0         0         0         0      Promoter (<=1kb)        23
#>          geneStart   geneEnd geneLength geneStrand      geneId      transcriptId
#>          <integer> <integer>  <integer>  <integer> <character>       <character>
#>    183 154429343 154449979      20637          1        3570 ENST00000476006.5
#>   1096   2181571   2186183       4613          1        6595 ENST00000635392.1
#>   2615  30292008  30307890      15883          2       83874 ENST00000403362.5
#>     34  10660737  10693912      33176          2       54897 ENST00000478728.2
#>    792  30743199  30744547       1349          2        8870 ENST00000259874.6
#>    ...       ...       ...        ...        ...         ...               ...
#>   2743 153920715 153926860       6146          2         393 ENST00000422091.1
#>   2745 154348524 154371203      22680          2        2316 ENST00000420627.5
#>   2746 154348529 154371283      22755          2        2316 ENST00000422373.6
#>   2747 154348529 154371283      22755          2        2316 ENST00000422373.6
#>   2748 154506204 154516242      10039          2       60343 ENST00000434658.6
#>          distanceToTSS         ENSEMBL      SYMBOL               GENENAME
#>              <numeric>     <character> <character>            <character>
#>    183           -9659 ENSG00000160712        IL6R interleukin 6 receptor
#>   1096           60798 ENSG00000080503     SMARCA2 SWI/SNF related, mat..
#>   2615           -2855 ENSG00000099992    TBC1D10A TBC1 domain family m..
#>     34            5242 ENSG00000130940       CASZ1   castor zinc finger 1
#>    792           -3891 ENSG00000137331        IER3 immediate early resp..
#>    ...             ...             ...         ...                    ...
#>   2743            -479 ENSG00000089820     ARHGAP4 Rho GTPase activatin..
#>   2745            1020 ENSG00000196924        FLNA              filamin A
#>   2746            -688 ENSG00000196924        FLNA              filamin A
#>   2747           -1067 ENSG00000196924        FLNA              filamin A
#>   2748            -897 ENSG00000071889       FAM3A FAM3 metabolism regu..
#>   -------
#>   seqinfo: 595 sequences (1 circular) from hg38 genome

## Evaluate calls on chromosome 1
```

```
head(as.data.frame(
  epiRomics_putative_enhanceosome_fantom@annotations
  )[as.data.frame(epiRomics_putative_enhanceosome_fantom@annotations)$seqnames
    == "chr1",])
#>     seqnames     start       end width strand foxa2 mafb nkx2_2 nkx6_1 pdx1
#> 183     chr1 154418514 154419684  1171      *     2    2      1      2    1
#> 34      chr1  10685395  10688670  3276      *     1    0      1      1    2
#> 24      chr1   8170112   8170857   746      *     1    1      1      1    1
#> 67      chr1  21638834  21639978  1145      *     1    1      1      1    1
#> 71      chr1  22414515  22414806   292      *     1    1      1      1    1
#> 256     chr1 205318810 205319660   851      *     1    1      2      1    0
#>     h2az ChIP_Hits                                  annotation geneChr
#> 183    2        10   Intron (ENST00000622330.4/3570, intron 1 of 6)       1
#> 34     2         7 Intron (ENST00000377022.8/54897, intron 4 of 20)       1
#> 24     1         6                              Distal Intergenic       1
#> 67     1         6  Intron (ENST00000290101.8/5909, intron 2 of 26)       1
#> 71     1         6                              Distal Intergenic       1
#> 256    1         6                               Promoter (2-3kb)       1
#>     geneStart   geneEnd geneLength geneStrand    geneId      transcriptId
#> 183 154429343 154449979      20637          1      3570 ENST00000476006.5
#> 34   10660737  10693912      33176          2     54897 ENST00000478728.2
#> 24    8201518   8215207      13690          1 102724539 ENST00000670361.1
#> 67   21596221  21651820      55600          2      5909 ENST00000471600.6
#> 71   22428838  22511763      82926          1      9923 ENST00000650433.1
#> 256 205302063 205321745      19683          2     81788 ENST00000367157.6
#>     distanceToTSS         ENSEMBL    SYMBOL
#> 183         -9659 ENSG00000160712      IL6R
#> 34           5242 ENSG00000130940     CASZ1
#> 24         -30661 ENSG00000227634 LINC01714
#> 67          11842 ENSG00000076864   RAP1GAP
#> 71         -14032 ENSG00000184677    ZBTB40
#> 256          2085 ENSG00000163545     NUAK2
#>                                     GENENAME
#> 183                    interleukin 6 receptor
#> 34                      castor zinc finger 1
#> 24  long intergenic non-protein coding RNA 1714
#> 67               RAP1 GTPase activating protein
#> 71       zinc finger and BTB domain containing 40
#> 256                     NUAK family kinase 2

## Find Index

which(names(epiRomics_putative_enhanceosome_fantom@annotations) == 183)
#> [1] 1
```

### Transcription factor decision trees

ChIP dataset repositories are quite sizeable for many organisms and cell types, with the expectation to only grow larger. Many different TFs binding to a putative enhancer region may not be that meaningful in the context of your biological question. A further step would be to ask whether there are co-TFs that pop up together, and whether this pattern varies across the functional annotation of the genome, i.e. does the combination of two TFs on enhanceosomes change on the gene body compared to distal intergenic regions?

```
plot(epiRomics_predictors(epiRomics_putative_enhanceosome_fantom))
```

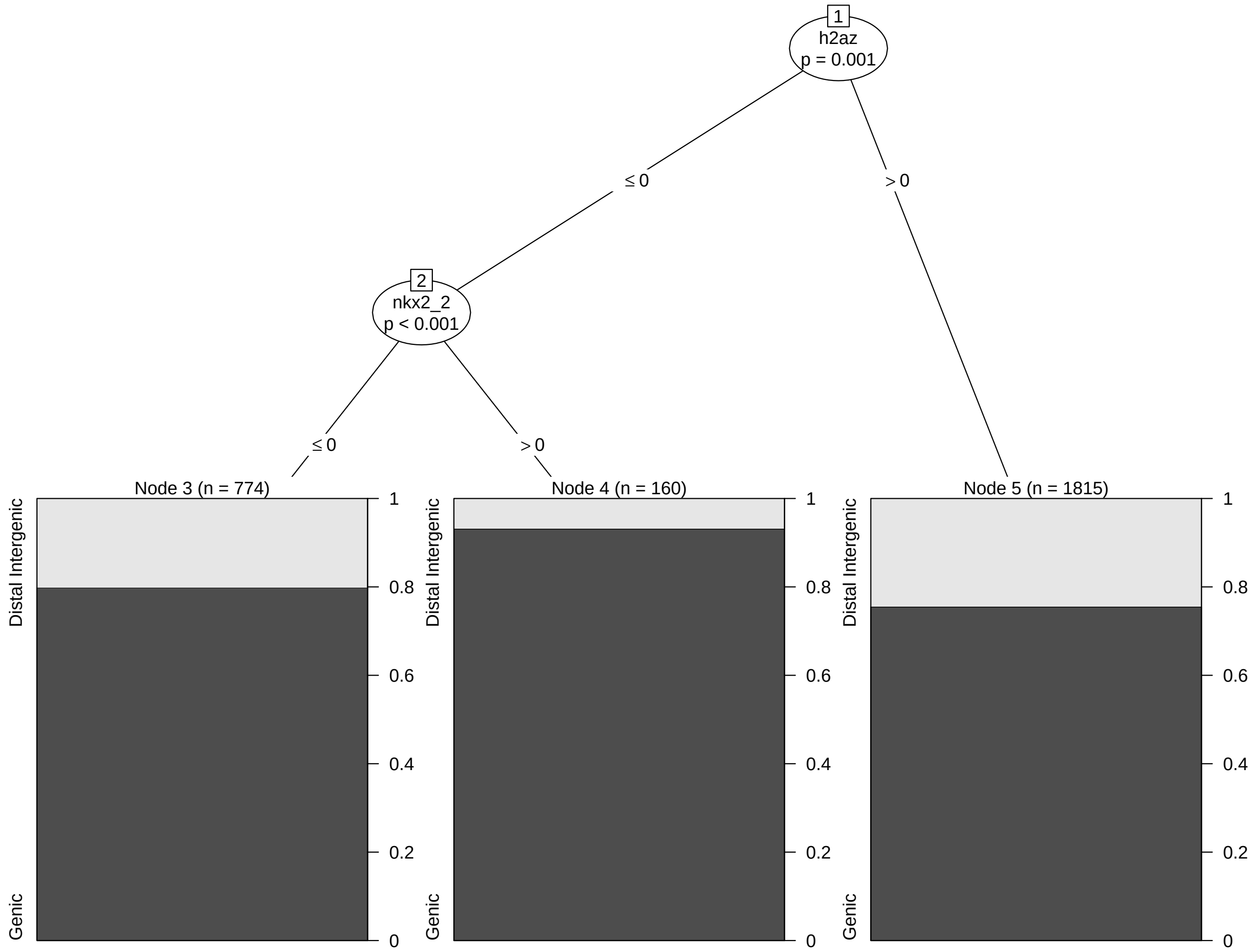

### Intersecting and visualizing ATAC- and RNA-Seq data

What if you wanted to visualize co-binding on your FANTOM filtered putative enhancer region? And do you have additional data you want to include for visualization, such as ATAC and RNA Seq? Lets take a look at one of the top hits

```
## Read in ATAC Seq and RNA Seq track bigwigs

## NOTE: These bigwigs are subsetted to chromosome 1.
## Indices not falling on chromosome 1 will return an error.

epiRomics_track_connection <- read.csv(
  system.file(
    "extdata",
    "example_epiRomics_BW_sheet_user_paths.csv",
    package = "epiRomics"
  )
)
```

```r
epiRomics_track_layer_human(
  epiRomics_putative_enhanceosome_fantom,
  epiRomics_index =
    which(
      names(epiRomics_putative_enhanceosome_fantom@annotations)
      == 183
    ),
  epiRomics_dB = epiRomics_dB,
  epiRomics_track_connection =
    epiRomics_track_connection
)
#> [1] "not empty"
#> [1] 103.678
#> [1] "not empty"
#> [1] 77.5726
#> [1] "not empty"
#> [1] 33.2945
#> [1] "not empty"
#> [1] 50.4959
```

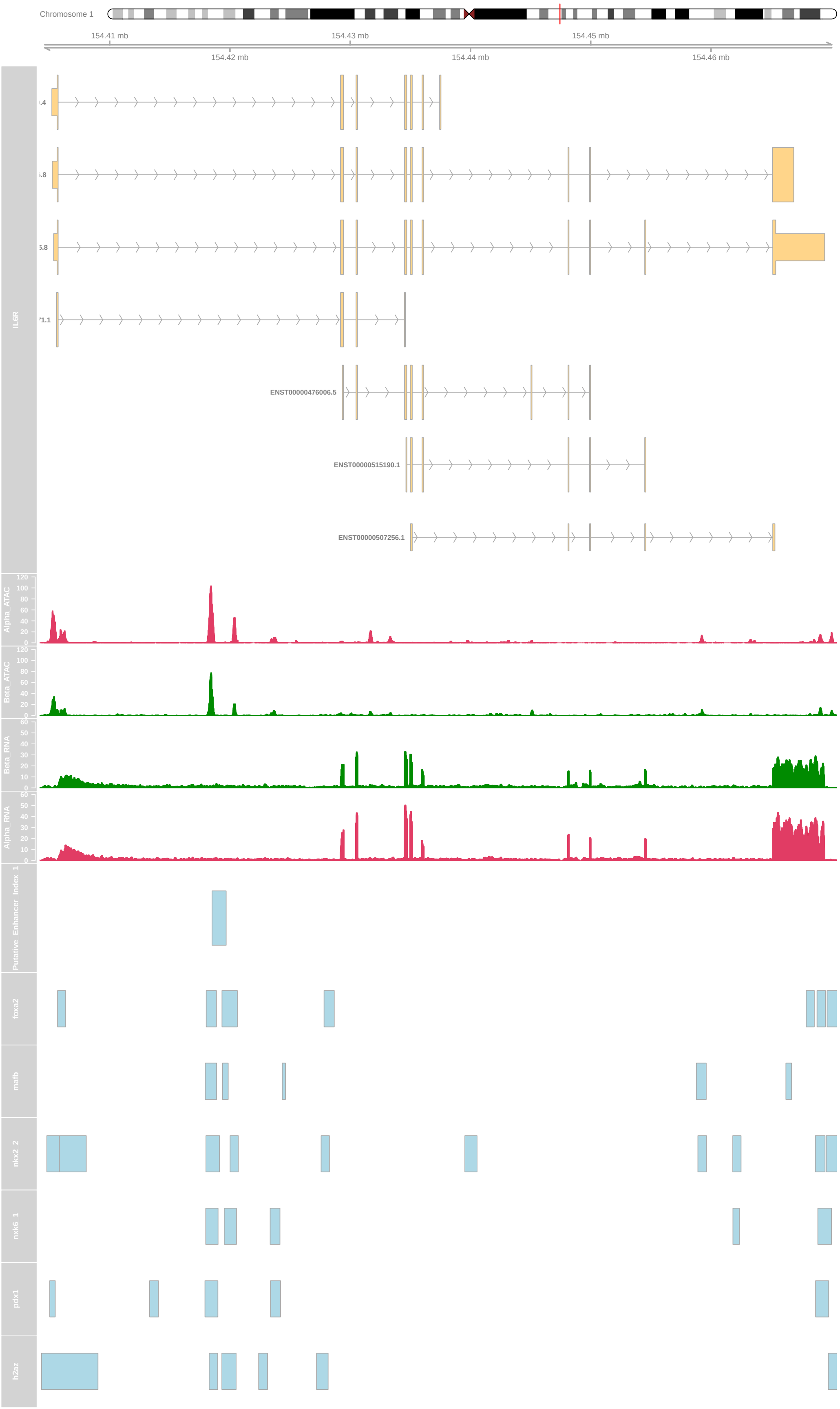
Chromosome 1
154.41 mb
154.43 mb
154.45 mb
154.42 mb
154.44 mb
154.46 mb
IL6R
ENST00000476006.5
ENST00000515190.1
ENST00000507256.1
Alpha_ATAC
Beta_ATAC
Beta_RNA
Alpha_RNA
Putative_Enhancer_Index_1
foxa2
mafb
nkx2_2
nkx6_1
pdx1
h2az

What about a region that overlapped with active enhancers from the human islet regulome database?

```
epiRomics_putative_enhanceosome_regulome_active <-
  epiRomics_enhanceosome(epiRomics_putative_enhancers_filtered_regulome_active,
                         epiRomics_dB)
#> >> preparing features information...      2021-10-05 17:46:34
#> >> identifying nearest features...        2021-10-05 17:46:34
#> >> calculating distance from peak to TSS...   2021-10-05 17:46:36
#> >> assigning genomic annotation...        2021-10-05 17:46:36
#> >> adding gene annotation...          2021-10-05 17:46:45
#> 'select()' returned 1:many mapping between keys and columns
#> >> assigning chromosome lengths          2021-10-05 17:46:45
#> >> done...                    2021-10-05 17:46:45

epiRomics_putative_enhanceosome_regulome_active@annotations
#> GRanges object with 6025 ranges and 19 metadata columns:
#>        seqnames              ranges strand |     foxa2      mafb    nkx2_2
#>           <Rle>           <IRanges>  <Rle> | <integer> <integer> <integer>
#>    456     chr1 154418514-154419684      * |         2         2         1
#>   2082     chr7     1555599-1556082      * |         1         1         1
#>   2572     chr9     2242369-2242873      * |         2         1         1
#>   3421    chr11   65416576-65419753      * |         1         1         2
#>   4709    chr17     7887867-7889135      * |         1         0         2
#>    ...      ...                 ...    ... .       ...       ...       ...
#>   5999     chrX            49184194      * |         0         0         0
#>   6001     chrX   70478965-70479351      * |         0         0         0
#>   6006     chrX 107710676-107711066      * |         0         0         0
#>   6007     chrX 107711430-107711673      * |         0         0         0
#>   6020     chrX 150874055-150874330      * |         0         0         0
#>           nkx6_1      pdx1      h2az ChIP_Hits             annotation   geneChr
#>        <integer> <integer> <integer> <numeric>            <character> <integer>
#>    456         2         1         2        10 Intron (ENST00000622..         1
#>   2082         2         2         1         8       Promoter (<=1kb)         7
#>   2572         1         2         1         8      Distal Intergenic         9
#>   3421         1         1         2         8      Distal Intergenic        11
#>   4709         2         2         1         8       Promoter (<=1kb)        17
#>    ...       ...       ...       ...       ...                    ...       ...
#>   5999         0         0         0         0       Promoter (<=1kb)        23
#>   6001         0         0         0         0       Promoter (<=1kb)        23
#>   6006         0         0         0         0      Distal Intergenic        23
#>   6007         0         0         0         0      Distal Intergenic        23
#>   6020         0         0         0         0 Intron (ENST00000370..        23
#>        geneStart   geneEnd geneLength geneStrand      geneId       transcriptId
#>        <integer> <integer>  <integer>  <integer> <character>        <character>
#>    456 154429343 154449979      20637          1        3570  ENST00000476006.5
#>   2082   1550305   1556120       5816          2      202915  ENST00000441933.5
#>   2572   2181571   2186183       4613          1        6595  ENST00000635392.1
#>   3421  65422798  65445540      22743          1      283131  ENST00000501122.2
#>   4709   7888789   7912755      23967          1        1107 ENST00000330494.12
#>    ...       ...       ...        ...        ...         ...                ...
#>   5999  49175621  49184789       9169          2        4007  ENST00000453382.5
#>   6001  70479118  70499903      20786          1        1741  ENST00000466140.5
#>   6006 107714677 107716401       1725          2        1831  ENST00000486554.1
```

```
#>   6007 107714677 107716401       1725          2        1831  ENST00000486554.1
#>   6020 150814900 150898609      83710          2       83692  ENST00000491877.1
#>          distanceToTSS         ENSEMBL      SYMBOL               GENENAME
#>              <numeric>     <character> <character>            <character>
#>    456           -9659 ENSG00000160712        IL6R interleukin 6 receptor
#>   2082              38 ENSG00000164855    TMEM184A transmembrane protei..
#>   2572           60798 ENSG00000080503     SMARCA2 SWI/SNF related, mat..
#>   3421           -3045 ENSG00000245532       NEAT1 nuclear paraspeckle ..
#>   4709               0 ENSG00000170004        CHD3 chromodomain helicas..
#>    ...             ...             ...         ...                    ...
#>   5999             595 ENSG00000012211    PRICKLE3 prickle planar cell ..
#>   6001               0 ENSG00000082458        DLG3 discs large MAGUK sc..
#>   6006            5335 ENSG00000157514     TSC22D3 TSC22 domain family ..
#>   6007            4728 ENSG00000157514     TSC22D3 TSC22 domain family ..
#>   6020           24279 ENSG00000102181      CD99L2   CD99 molecule like 2
#>   -------
#>   seqinfo: 595 sequences (1 circular) from hg38 genome

## Evaluate calls on chromosome 1

head(as.data.frame(
  epiRomics_putative_enhanceosome_regulome_active@annotations
)[as.data.frame(epiRomics_putative_enhanceosome_regulome_active@annotations)$seqnames
  == "chr1",])
#>     seqnames     start       end width strand foxa2 mafb nkx2_2 nkx6_1 pdx1
#> 456     chr1 154418514 154419684  1171      *     2    2      1      2    1
#> 82      chr1  10685395  10688670  3276      *     1    0      1      1    2
#> 46      chr1   7574092   7574479   388      *     1    1      1      1    1
#> 47      chr1   7574640   7575094   455      *     1    1      1      1    1
#> 49      chr1   8169274   8169689   416      *     1    1      1      1    1
#> 50      chr1   8170112   8170857   746      *     1    1      1      1    1
#>     h2az ChIP_Hits                                       annotation geneChr
#> 456    2        10    Intron (ENST00000622330.4/3570, intron 1 of 6)       1
#> 82     2         7  Intron (ENST00000377022.8/54897, intron 4 of 20)       1
#> 46     1         6 Intron (ENST00000303635.12/23261, intron 6 of 22)       1
#> 47     1         6 Intron (ENST00000303635.12/23261, intron 6 of 22)       1
#> 49     1         6                                Distal Intergenic       1
#> 50     1         6                                Distal Intergenic       1
#>     geneStart   geneEnd geneLength geneStrand    geneId      transcriptId
#> 456 154429343 154449979      20637          1      3570 ENST00000476006.5
#> 82   10660737  10693912      33176          2     54897 ENST00000478728.2
#> 46    7736408   7767856      31449          1     23261 ENST00000495233.5
#> 47    7736408   7767856      31449          1     23261 ENST00000495233.5
#> 49    8201518   8215207      13690          1 102724539 ENST00000670361.1
#> 50    8201518   8215207      13690          1 102724539 ENST00000670361.1
#>     distanceToTSS         ENSEMBL    SYMBOL
#> 456         -9659 ENSG00000160712      IL6R
#> 82           5242 ENSG00000130940     CASZ1
#> 46        -161929 ENSG00000171735    CAMTA1
#> 47        -161314 ENSG00000171735    CAMTA1
#> 49         -31829 ENSG00000227634 LINC01714
#> 50         -30661 ENSG00000227634 LINC01714
#>                                      GENENAME
```

```r
#> 456                       interleukin 6 receptor
#> 82                          castor zinc finger 1
#> 46  calmodulin binding transcription activator 1
#> 47  calmodulin binding transcription activator 1
#> 49   long intergenic non-protein coding RNA 1714
#> 50   long intergenic non-protein coding RNA 1714

## Find Index

which(names(epiRomics_putative_enhanceosome_regulome_active@annotations) == 82)
#> [1] 7
```

```r
epiRomics_track_layer_human(
  epiRomics_putative_enhanceosome_regulome_active,
  epiRomics_index = which(
    names(
      epiRomics_putative_enhanceosome_regulome_active@annotations
    ) == 82
  ),
  epiRomics_dB = epiRomics_dB,
  epiRomics_track_connection = epiRomics_track_connection
)
#> [1] "not empty"
#> [1] 36.1883
#> [1] "not empty"
#> [1] 14.7321
#> [1] "not empty"
#> [1] 5.30505
#> [1] "not empty"
#> [1] 2.40628
```

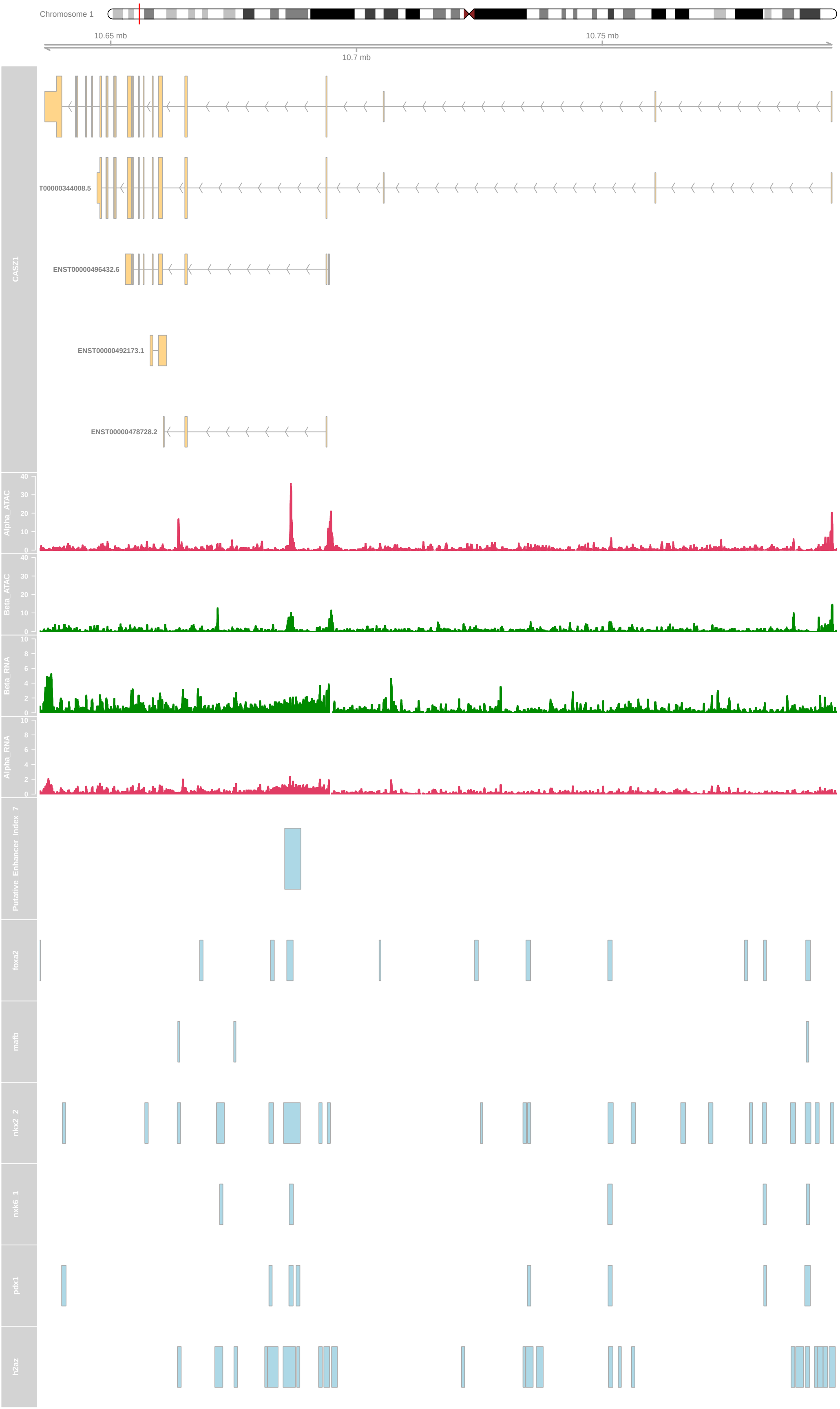

Chromosome 1
10.65 mb
10.7 mb
10.75 mb
CASZ1
T00000344008.5
ENST00000496432.6
ENST00000492173.1
ENST00000478728.2
Alpha_ATAC
Beta_ATAC
Beta_RNA
Alpha_RNA
Putative_Enhancer_Index_7
foxa2
mafb
nkx2_2
nkx6_1
pdx1
h2az

What about a region that overlapped with super enhancers from the human islet regulome database?

```
epiRomics_putative_enhanceosome_regulome_super <-
  epiRomics_enhanceosome(epiRomics_putative_enhancers_filtered_regulome_super,
                         epiRomics_dB)
#> >> preparing features information...      2021-10-05 17:49:52
#> >> identifying nearest features...        2021-10-05 17:49:52
#> >> calculating distance from peak to TSS...   2021-10-05 17:49:53
#> >> assigning genomic annotation...        2021-10-05 17:49:53
#> >> adding gene annotation...          2021-10-05 17:50:00
#> 'select()' returned 1:many mapping between keys and columns
#> >> assigning chromosome lengths           2021-10-05 17:50:01
#> >> done...                   2021-10-05 17:50:01

epiRomics_putative_enhanceosome_regulome_super@annotations
#> GRanges object with 2401 ranges and 19 metadata columns:
#>          seqnames              ranges strand |     foxa2      mafb    nkx2_2
#>             <Rle>           <IRanges>  <Rle> | <integer> <integer> <integer>
#>    166      chr1 154418514-154419684      * |         2         2         1
#>    966      chr9     2242369-2242873      * |         2         1         1
#>   1422     chr11   65416576-65419753      * |         1         1         2
#>     16      chr1   10685395-10688670      * |         1         0         1
#>    764      chr6   30748438-30749427      * |         2         1         1
#>    ...       ...                 ...    ... .       ...       ...       ...
#>   2393     chr22   46103687-46105878      * |         0         0         0
#>   2396     chr22   46108912-46109315      * |         0         0         0
#>   2397     chr22   46109916-46110442      * |         0         0         0
#>   2398     chr22   46115774-46116154      * |         0         0         0
#>   2399     chr22   46116326-46116501      * |         0         0         0
#>             nkx6_1      pdx1      h2az ChIP_Hits             annotation   geneChr
#>          <integer> <integer> <integer> <numeric>            <character> <integer>
#>    166          2         1         2        10 Intron (ENST00000622..         1
#>    966          1         2         1         8      Distal Intergenic         9
#>   1422          1         1         2         8      Distal Intergenic        11
#>     16          1         2         2         7 Intron (ENST00000377..         1
#>    764          1         1         1         7 Intron (ENST00000656..         6
#>    ...        ...       ...       ...       ...                    ...       ...
#>   2393          0         0         0         0 Exon (ENST0000038105..        22
#>   2396          0         0         0         0 Exon (ENST0000043543..        22
#>   2397          0         0         0         0       Promoter (2-3kb)        22
#>   2398          0         0         0         0       Promoter (2-3kb)        22
#>   2399          0         0         0         0       Promoter (2-3kb)        22
#>          geneStart   geneEnd geneLength geneStrand      geneId      transcriptId
#>          <integer> <integer>  <integer>  <integer> <character>       <character>
#>    166 154429343 154449979      20637          1        3570 ENST00000476006.5
#>    966   2181571   2186183       4613          1        6595 ENST00000635392.1
#>   1422  65422798  65445540      22743          1      283131 ENST00000501122.2
#>     16  10660737  10693912      33176          2       54897 ENST00000478728.2
#>    764  30743199  30744547       1349          2        8870 ENST00000259874.6
#>    ...       ...       ...        ...        ...         ...               ...
#>   2393  46112749  46112822         74          1      406883 ENST00000362116.3
#>   2396  46112749  46112822         74          1      406883 ENST00000362116.3
#>   2397  46112749  46112822         74          1      406883 ENST00000362116.3
```

```
#>   2398  46113686  46113768         83          1      406884 ENST00000385140.1
#>   2399  46113686  46113768         83          1      406884 ENST00000385140.1
#>        distanceToTSS         ENSEMBL      SYMBOL               GENENAME
#>            <numeric>     <character> <character>            <character>
#>    166         -9659 ENSG00000160712        IL6R interleukin 6 receptor
#>    966         60798 ENSG00000080503     SMARCA2 SWI/SNF related, mat..
#>   1422         -3045 ENSG00000245532       NEAT1 nuclear paraspeckle ..
#>     16          5242 ENSG00000130940       CASZ1   castor zinc finger 1
#>    764         -3891 ENSG00000137331        IER3 immediate early resp..
#>    ...           ...             ...         ...                    ...
#>   2393         -6871 ENSG00000283990    MIRLET7A3      microRNA let-7a-3
#>   2396         -3434 ENSG00000283990    MIRLET7A3      microRNA let-7a-3
#>   2397         -2307 ENSG00000283990    MIRLET7A3      microRNA let-7a-3
#>   2398          2088 ENSG00000284520     MIRLET7B        microRNA let-7b
#>   2399          2640 ENSG00000284520     MIRLET7B        microRNA let-7b
#>   -------
#>   seqinfo: 595 sequences (1 circular) from hg38 genome

## Evaluate calls on chromosome 1

head(as.data.frame(
  epiRomics_putative_enhanceosome_regulome_super@annotations)[as.data.frame(
    epiRomics_putative_enhanceosome_regulome_super@annotations)$seqnames == "chr1",])
#>     seqnames     start       end width strand foxa2 mafb nkx2_2 nkx6_1 pdx1
#> 166     chr1 154418514 154419684  1171      *     2    2      1      2    1
#> 16      chr1  10685395  10688670  3276      *     1    0      1      1    2
#> 1       chr1   7574092   7574479   388      *     1    1      1      1    1
#> 2       chr1   7574640   7575094   455      *     1    1      1      1    1
#> 3       chr1   8169274   8169689   416      *     1    1      1      1    1
#> 4       chr1   8170112   8170857   746      *     1    1      1      1    1
#>     h2az ChIP_Hits                                      annotation geneChr
#> 166    2        10    Intron (ENST00000622330.4/3570, intron 1 of 6)       1
#> 16     2         7  Intron (ENST00000377022.8/54897, intron 4 of 20)       1
#> 1      1         6 Intron (ENST00000303635.12/23261, intron 6 of 22)       1
#> 2      1         6 Intron (ENST00000303635.12/23261, intron 6 of 22)       1
#> 3      1         6                               Distal Intergenic       1
#> 4      1         6                               Distal Intergenic       1
#>     geneStart   geneEnd geneLength geneStrand    geneId      transcriptId
#> 166 154429343 154449979      20637          1      3570 ENST00000476006.5
#> 16   10660737  10693912      33176          2     54897 ENST00000478728.2
#> 1     7736408   7767856      31449          1     23261 ENST00000495233.5
#> 2     7736408   7767856      31449          1     23261 ENST00000495233.5
#> 3     8201518   8215207      13690          1 102724539 ENST00000670361.1
#> 4     8201518   8215207      13690          1 102724539 ENST00000670361.1
#>     distanceToTSS         ENSEMBL    SYMBOL
#> 166         -9659 ENSG00000160712      IL6R
#> 16           5242 ENSG00000130940     CASZ1
#> 1         -161929 ENSG00000171735    CAMTA1
#> 2         -161314 ENSG00000171735    CAMTA1
#> 3          -31829 ENSG00000227634 LINC01714
#> 4          -30661 ENSG00000227634 LINC01714
#>                                  GENENAME
#> 166                 interleukin 6 receptor
```

```r
#> 16                          castor zinc finger 1
#> 1   calmodulin binding transcription activator 1
#> 2   calmodulin binding transcription activator 1
#> 3    long intergenic non-protein coding RNA 1714
#> 4    long intergenic non-protein coding RNA 1714

## Find Index

which(names(epiRomics_putative_enhanceosome_regulome_super@annotations) == 1)
#> [1] 14
```

```r
epiRomics_track_layer_human(
  epiRomics_putative_enhanceosome_regulome_super,
  epiRomics_index = which(
    names(epiRomics_putative_enhanceosome_regulome_super@annotations) == 1
  ),
  epiRomics_dB = epiRomics_dB,
  epiRomics_track_connection = epiRomics_track_connection
)
#> [1] "not empty"
#> [1] 243.743
#> [1] "not empty"
#> [1] 103.706
#> [1] "not empty"
#> [1] 11.3213
#> [1] "not empty"
#> [1] 22.1498
```

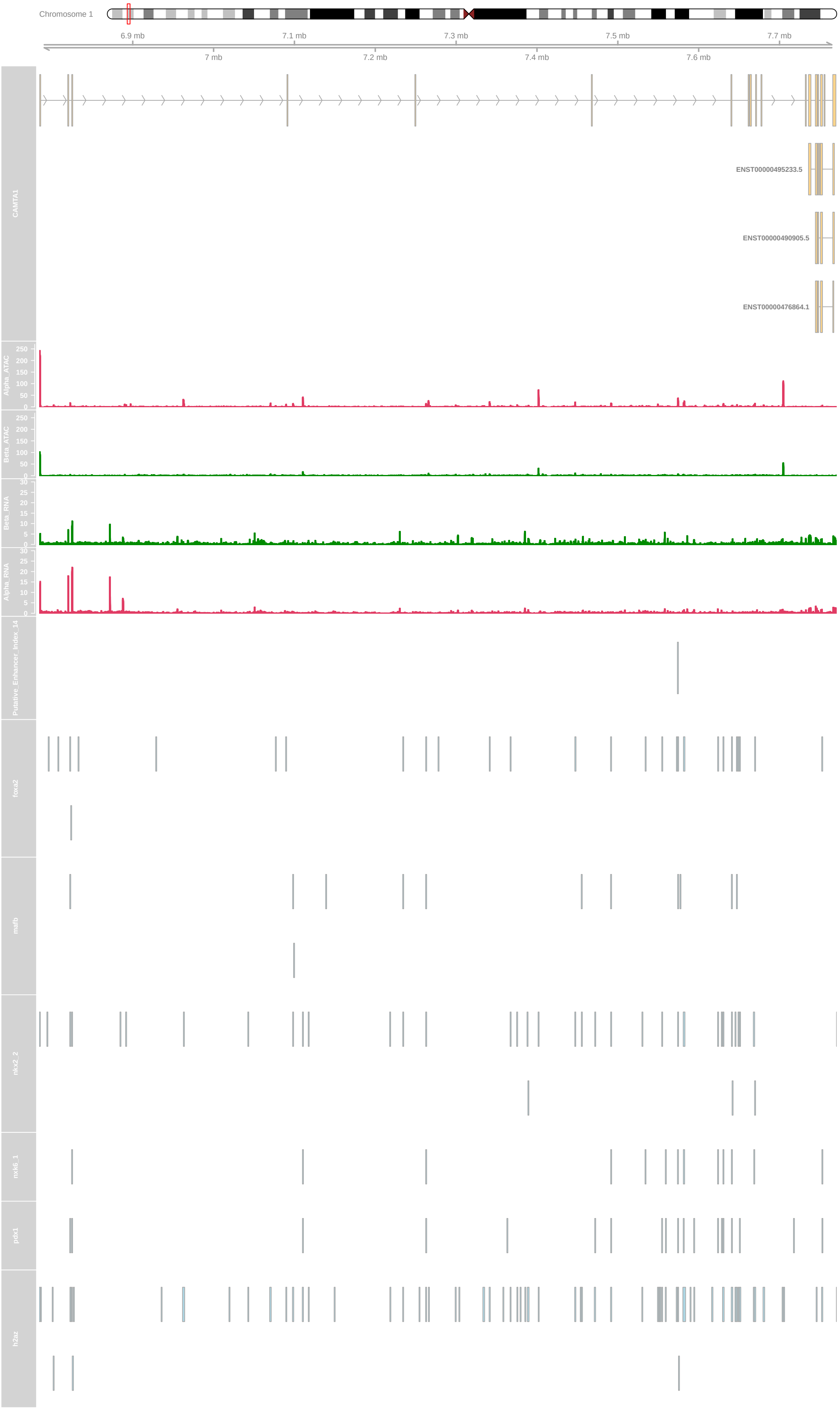
Chromosome 1
6.9 mb
7.1 mb
7.3 mb
7.5 mb
7.7 mb
7 mb
7.2 mb
7.4 mb
7.6 mb
CAMTA1
ENST00000495233.5
ENST00000490905.5
ENST00000476864.1
Alpha_ATAC
Beta_ATAC
Beta_RNA
Alpha_RNA
Putative_Enhancer_Index_14
foxa2
mafb
nkx2_2
nkx6_1
pdx1
h2az

Or, about a region that overlapped with ultra-conserved non coding elements?

```
epiRomics_putative_enhanceosome_ucnes <-
  epiRomics_enhanceosome(epiRomics_putative_enhancers_filtered_ucnes, epiRomics_dB)
#> >> preparing features information...      2021-10-05 17:52:00
#> >> identifying nearest features...        2021-10-05 17:52:00
#> >> calculating distance from peak to TSS...   2021-10-05 17:52:01
#> >> assigning genomic annotation...        2021-10-05 17:52:01
#> >> adding gene annotation...          2021-10-05 17:52:08
#> 'select()' returned 1:1 mapping between keys and columns
#> >> assigning chromosome lengths          2021-10-05 17:52:08
#> >> done...                    2021-10-05 17:52:08

epiRomics_putative_enhanceosome_ucnes@annotations
#> GRanges object with 11 ranges and 19 metadata columns:
#>      seqnames              ranges strand |     foxa2      mafb    nkx2_2
#>         <Rle>           <IRanges>  <Rle> | <integer> <integer> <integer>
#>    7     chr9 106921420-106921764      * |         1         1         1
#>    1     chr1 164635220-164635921      * |         0         0         2
#>    6     chr3   71131859-71132164      * |         1         0         0
#>    2     chr1 164711914-164712296      * |         0         0         0
#>    3     chr1 164712350-164713071      * |         0         0         0
#>    8    chr11 114163425-114164860      * |         0         0         0
#>    9    chr15   36903894-36904085      * |         0         0         1
#>   11    chr21   16534340-16534665      * |         0         0         0
#>    4     chr1 200079185-200079426      * |         0         0         0
#>    5     chr1 213585694-213586385      * |         0         0         0
#>   10    chr15   53447393-53447809      * |         0         0         0
#>         nkx6_1      pdx1      h2az ChIP_Hits             annotation   geneChr
#>      <integer> <integer> <integer> <numeric>            <character> <integer>
#>    7         1         0         1         5 Intron (ENST00000472..         9
#>    1         0         1         1         4 Intron (ENST00000420..         1
#>    6         0         0         1         2       Promoter (<=1kb)         3
#>    2         0         0         1         1 Intron (ENST00000420..         1
#>    3         0         0         1         1 Intron (ENST00000420..         1
#>    8         0         0         1         1 Intron (ENST00000335..        11
#>    9         0         0         0         1       Promoter (<=1kb)        15
#>   11         0         0         1         1       Promoter (<=1kb)        21
#>    4         0         0         0         0 Intron (ENST00000236..         1
#>    5         0         0         0         0      Distal Intergenic         1
#>   10         0         0         0         0 Intron (ENST00000662..        15
#>      geneStart   geneEnd geneLength geneStrand      geneId      transcriptId
#>      <integer> <integer>  <integer>  <integer> <character>       <character>
#>    7 106926925 106932462       5538          1       58499 ENST00000480607.5
#>    1 164630981 164799889     168909          1        5087 ENST00000482110.5
#>    6  70959237  71132099     172863          2       27086 ENST00000650188.1
#>    2 164772912 164807571      34660          1        5087 ENST00000558837.5
#>    3 164772912 164807571      34660          1        5087 ENST00000558837.5
#>    8 114180766 114247296      66531          1        7704 ENST00000545851.5
#>    9  36894784  36904067       9284          2        4212 ENST00000559408.1
#>   11  16534952  16607137      72186          1      388815 ENST00000654245.1
#>    4 200043810 200058424      14615          1        2494 ENST00000367357.3
#>    5 213832591 213841041       8451          2   100505832 ENST00000609394.5
```

```
#>   10  53513742  53541080      27339          2      256764 ENST00000614174.4
#>      distanceToTSS         ENSEMBL      SYMBOL               GENENAME
#>          <numeric>     <character> <character>            <character>
#>    7         -5161 ENSG00000148143      ZNF462 zinc finger protein ..
#>    1          4239 ENSG00000185630        PBX1          PBX homeobox 1
#>    6             0 ENSG00000114861       FOXP1         forkhead box P1
#>    2        -60616 ENSG00000185630        PBX1          PBX homeobox 1
#>    3        -59841 ENSG00000185630        PBX1          PBX homeobox 1
#>    8        -15906 ENSG00000109906      ZBTB16 zinc finger and BTB ..
#>    9             0 ENSG00000134138       MEIS2         Meis homeobox 2
#>   11          -287 ENSG00000215386     MIR99AHG mir-99a-let-7c clust..
#>    4         35375 ENSG00000116833       NR5A2 nuclear receptor sub..
#>    5        254656 ENSG00000230461    PROX1-AS1  PROX1 antisense RNA 1
#>   10         93271 ENSG00000166415       WDR72     WD repeat domain 72
#>   -------
#>   seqinfo: 595 sequences (1 circular) from hg38 genome
```

```
epiRomics_track_layer_human(
  epiRomics_putative_enhanceosome_ucnes,
  epiRomics_index = 9,
  epiRomics_dB = epiRomics_dB,
  epiRomics_track_connection = epiRomics_track_connection
)
#> [1] "not empty"
#> [1] 82.795
#> [1] "not empty"
#> [1] 62.847
#> [1] "not empty"
#> [1] 3.55117
#> [1] "not empty"
#> [1] 3.80884
```

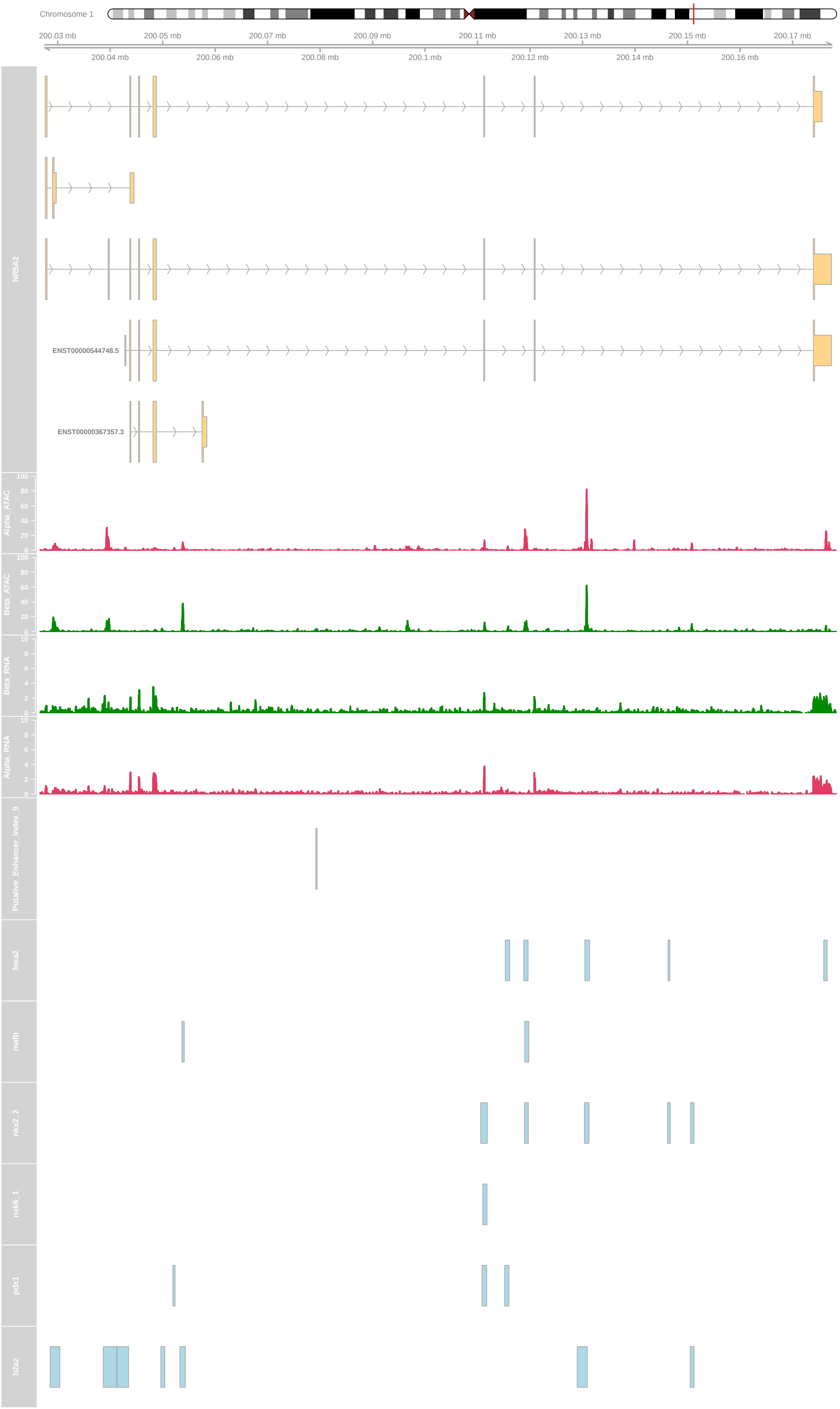
Chromosome 1
200.03 mb
200.04 mb
200.05 mb
200.06 mb
200.07 mb
200.08 mb
200.09 mb
200.1 mb
200.11 mb
200.12 mb
200.13 mb
200.14 mb
200.15 mb
200.16 mb
200.17 mb
NR5A2
ENST00000544748.5
ENST00000367357.3
Alpha_ATAC
Beta_ATAC
Beta_RNA
Alpha_RNA
Putative_Enhancer_Index_9
foxa2
mafb
nkx2_2
nkx6_1
pdx1
h2az

How about applying multiple filters to further increase the confidence of calls?

```
epiRomics_putative_enhancers_filtered_stringent <-
  epiRomics_enhancers_filter(
    epiRomics_enhancers_filter(
      epiRomics_enhancers_filter(
        epiRomics_enhancers_filter(epiRomics_putative_enhancers, epiRomics_dB,
                                   epiRomics_type =
                                     "hg38_custom_fantom"),
        epiRomics_dB,
        epiRomics_type = "hg38_custom_regulome_active"
      ),
      epiRomics_dB,
      epiRomics_type = "hg38_custom_regulome_super"
    ),
    epiRomics_dB,
    epiRomics_type = "hg38_custom_ucnes"
  )

## Here, we see a highly conservative list of putative enhancer calls that overlap with four
## different functional annotations, suggesting the lowest hanging fruit for downstream
## bench-lab validation.
## NOTE: The UCNE database filter caused the greatest reduction in enhancer calls.

epiRomics_putative_enhancers_filtered_stringent@annotations
#> GRanges object with 2 ranges and 0 metadata columns:
#>       seqnames              ranges strand
#>          <Rle>           <IRanges>  <Rle>
#>   [1]     chr1 164711914-164712296      *
#>   [2]     chr1 164712350-164713071      *
#>   -------
#>   seqinfo: 595 sequences (1 circular) from hg38 genome

epiRomics_putative_enhanceosome_stringent <-
  epiRomics_enhanceosome(epiRomics_putative_enhancers_filtered_stringent,
                         epiRomics_dB)
#> >> preparing features information...      2021-10-05 17:54:01
#> >> identifying nearest features...        2021-10-05 17:54:01
#> >> calculating distance from peak to TSS...   2021-10-05 17:54:02
#> >> assigning genomic annotation...        2021-10-05 17:54:02
#> >> adding gene annotation...          2021-10-05 17:54:09
#> 'select()' returned 1:1 mapping between keys and columns
#> >> assigning chromosome lengths           2021-10-05 17:54:09
#> >> done...                    2021-10-05 17:54:09
```

```
epiRomics_track_layer_human(
  epiRomics_putative_enhanceosome_stringent,
  epiRomics_index = 1,
  epiRomics_dB = epiRomics_dB,
  epiRomics_track_connection = epiRomics_track_connection
)
#> [1] "not empty"
```

```
#> [1] 233.705
#> [1] "not empty"
#> [1] 221.083
#> [1] "not empty"
#> [1] 9.31751
#> [1] "not empty"
#> [1] 15.8259
```

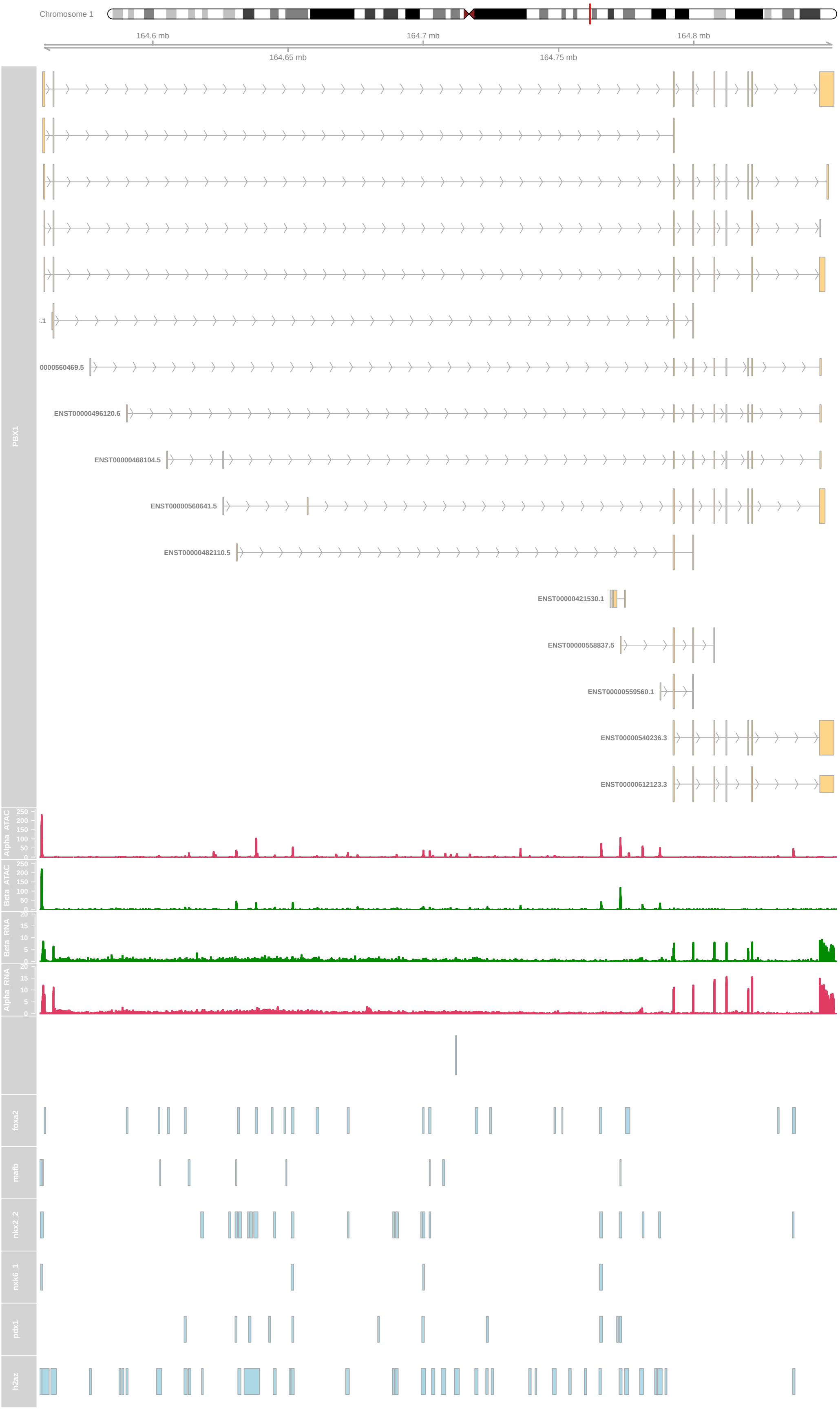
Chromosome 1
164.6 mb
164.7 mb
164.8 mb
164.65 mb
164.75 mb
PBX1
0000560469.5
ENST00000496120.6
ENST00000468104.5
ENST00000560641.5
ENST00000482110.5
ENST00000421530.1
ENST00000558837.5
ENST00000559560.1
ENST00000540236.3
ENST00000612123.3
Alpha_ATAC
Beta_ATAC
Beta_RNA
Alpha_RNA
foxa2
mafb
nkx2_2
nkx6_1
pdx1
h2az

How can we use these putative enhanceosome regions to infer biology between cell states? In this example, we will integrate ATAC-Seq data differential testing showing differences in chromatin accessibility between alpha and beta cells

```
## Read differentially binding data generated with DiffBind.
## DE comparing human alpha and beta cell chromatin.

b.v.a <-
  read.csv(system.file("extdata", "DBA_Beta_Versus_Alpha.csv", package = "epiRomics"))
b.v.a <- GRanges(b.v.a)

# Filter for beta enriched chromatin regions

beta.enriched <- b.v.a[b.v.a$Fold >= 1, ]

# Connect to our putative enhanceosomes

beta_enhancer_regions <-
  epiRomics_regions_of_interest(epiRomics_putative_enhanceosome_fantom, beta.enriched)
```

Now, lets visualize the top candidate region we found after connecting our differential chromatin analysis with the putative enhanceosomes

```
epiRomics_track_layer_human(
  beta_enhancer_regions,
  epiRomics_index = 1,
  epiRomics_dB = epiRomics_dB,
  epiRomics_track_connection = epiRomics_track_connection
)
#> [1] "not empty"
#> [1] 16.5139
#> [1] "not empty"
#> [1] 14.2229
#> [1] "not empty"
#> [1] 2.38379
#> [1] "not empty"
#> [1] 1.29041
```

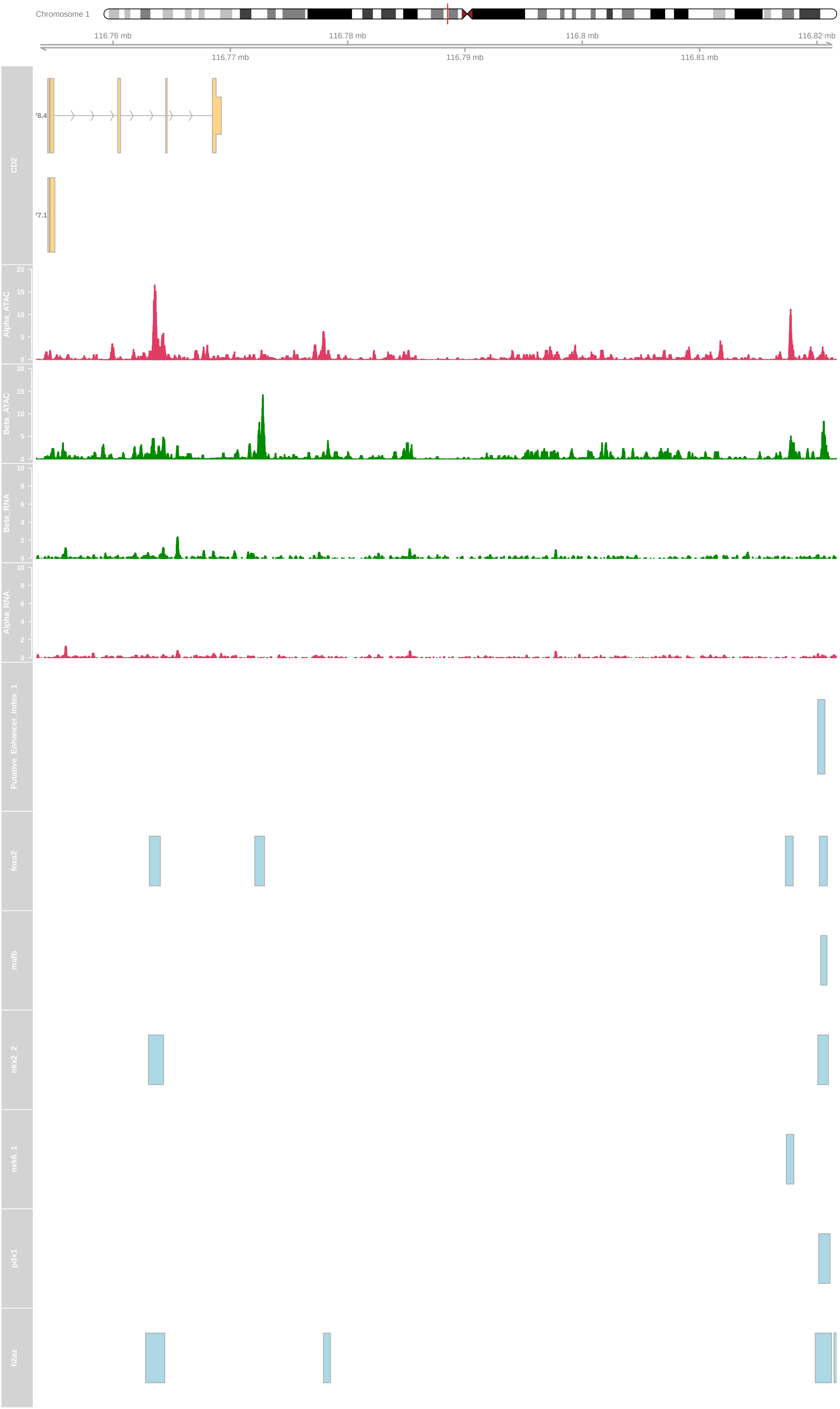
Chromosome 1
116.76 mb
116.78 mb
116.8 mb
116.82 mb
116.77 mb
116.79 mb
116.81 mb
CD2
Alpha_ATAC
Beta_ATAC
Beta_RNA
Alpha_RNA
Putative_Enhancer_Index_1
foxa2
mafb
nkx2_2
nkx6_1
pdx1
h2az

#### Session Information

Here is the output of `sessionInfo()` on the system on which this document was compiled:

```
sessionInfo()
#> R version 4.1.1 (2021-08-10)
#> Platform: x86_64-apple-darwin17.0 (64-bit)
#> Running under: macOS Catalina 10.15.7
#>
#> Matrix products: default
#> BLAS:   /Library/Frameworks/R.framework/Versions/4.1/Resources/lib/libRblas.0.dylib
#> LAPACK: /Library/Frameworks/R.framework/Versions/4.1/Resources/lib/libRlapack.dylib
#>
#> locale:
#> [1] en_US.UTF-8/en_US.UTF-8/en_US.UTF-8/C/en_US.UTF-8/en_US.UTF-8
#>
#> attached base packages:
#> [1] stats4    parallel  stats     graphics  grDevices utils     datasets
#> [8] methods   base
#>
#> other attached packages:
#>  [1] BSgenome.Hsapiens.UCSC.hg38_1.4.3
#>  [2] BSgenome_1.60.0
#>  [3] rtracklayer_1.52.1
#>  [4] Biostrings_2.60.2
#>  [5] XVector_0.32.0
#>  [6] org.Hs.eg.db_3.13.0
#>  [7] TxDb.Hsapiens.UCSC.hg38.knownGene_3.13.0
#>  [8] GenomicFeatures_1.44.2
#>  [9] AnnotationDbi_1.54.1
#> [10] Biobase_2.52.0
#> [11] GenomicRanges_1.44.0
#> [12] GenomeInfoDb_1.28.4
#> [13] IRanges_2.26.0
#> [14] S4Vectors_0.30.2
#> [15] BiocGenerics_0.38.0
#> [16] epiRomics_0.1.3
#>
#> loaded via a namespace (and not attached):
#>   [1] utf8_1.2.2
#>   [2] tidyselect_1.1.1
#>   [3] RSQLite_2.2.8
#>   [4] htmlwidgets_1.5.4
#>   [5] grid_4.1.1
#>   [6] BiocParallel_1.26.2
#>   [7] scatterpie_0.1.7
#>   [8] munsell_0.5.0
#>   [9] codetools_0.2-18
#>  [10] withr_2.4.2
#>  [11] colorspace_2.0-2
#>  [12] GOSemSim_2.18.1
#>  [13] filelock_1.0.2
#>  [14] knitr_1.36
```

```
#>  [15] rstudioapi_0.13
#>  [16] DOSE_3.18.3
#>  [17] MatrixGenerics_1.4.3
#>  [18] GenomeInfoDbData_1.2.6
#>  [19] polyclip_1.10-0
#>  [20] bit64_4.0.5
#>  [21] farver_2.1.0
#>  [22] treeio_1.16.2
#>  [23] vctrs_0.3.8
#>  [24] generics_0.1.0
#>  [25] TH.data_1.1-0
#>  [26] xfun_0.26
#>  [27] biovizBase_1.40.0
#>  [28] BiocFileCache_2.0.0
#>  [29] party_1.3-9
#>  [30] regioneR_1.24.0
#>  [31] R6_2.5.1
#>  [32] graphlayouts_0.7.1
#>  [33] AnnotationFilter_1.16.0
#>  [34] bitops_1.0-7
#>  [35] cachem_1.0.6
#>  [36] fgsea_1.18.0
#>  [37] gridGraphics_0.5-1
#>  [38] DelayedArray_0.18.0
#>  [39] assertthat_0.2.1
#>  [40] vroom_1.5.5
#>  [41] promises_1.2.0.1
#>  [42] BiocIO_1.2.0
#>  [43] scales_1.1.1
#>  [44] multcomp_1.4-17
#>  [45] ggraph_2.0.5
#>  [46] nnet_7.3-16
#>  [47] enrichplot_1.13.1.992
#>  [48] gtable_0.3.0
#>  [49] tidygraph_1.2.0
#>  [50] sandwich_3.0-1
#>  [51] ensembldb_2.16.4
#>  [52] rlang_0.4.11
#>  [53] splines_4.1.1
#>  [54] lazyeval_0.2.2
#>  [55] dichromat_2.0-0
#>  [56] checkmate_2.0.0
#>  [57] BiocManager_1.30.16
#>  [58] yaml_2.2.1
#>  [59] reshape2_1.4.4
#>  [60] backports_1.2.1
#>  [61] httpuv_1.6.3
#>  [62] qvalue_2.24.0
#>  [63] Hmisc_4.5-0
#>  [64] tools_4.1.1
#>  [65] ggplotify_0.1.0
#>  [66] ggplot2_3.3.5
#>  [67] gplots_3.1.1
```

```
#>  [68] ellipsis_0.3.2
#>  [69] RColorBrewer_1.1-2
#>  [70] Rcpp_1.0.7
#>  [71] plyr_1.8.6
#>  [72] base64enc_0.1-3
#>  [73] progress_1.2.2
#>  [74] zlibbioc_1.38.0
#>  [75] purrr_0.3.4
#>  [76] RCurl_1.98-1.5
#>  [77] prettyunits_1.1.1
#>  [78] rpart_4.1-15
#>  [79] viridis_0.6.1
#>  [80] zoo_1.8-9
#>  [81] SummarizedExperiment_1.22.0
#>  [82] ggrepel_0.9.1
#>  [83] cluster_2.1.2
#>  [84] magrittr_2.0.1
#>  [85] data.table_1.14.2
#>  [86] DO.db_2.9
#>  [87] mvtnorm_1.1-2
#>  [88] ProtGenerics_1.24.0
#>  [89] matrixStats_0.61.0
#>  [90] hms_1.1.1
#>  [91] patchwork_1.1.1
#>  [92] mime_0.12
#>  [93] evaluate_0.14
#>  [94] xtable_1.8-4
#>  [95] XML_3.99-0.8
#>  [96] jpeg_0.1-9
#>  [97] gridExtra_2.3
#>  [98] compiler_4.1.1
#>  [99] biomaRt_2.48.3
#> [100] tibble_3.1.5
#> [101] KernSmooth_2.23-20
#> [102] shadowtext_0.0.9
#> [103] crayon_1.4.1
#> [104] htmltools_0.5.2
#> [105] ggfun_0.0.4
#> [106] later_1.3.0
#> [107] tzdb_0.1.2
#> [108] Formula_1.2-4
#> [109] tidyr_1.1.4
#> [110] aplot_0.1.1
#> [111] libcoin_1.0-9
#> [112] DBI_1.1.1
#> [113] formatR_1.11
#> [114] ChIPseeker_1.28.3
#> [115] tweenr_1.0.2
#> [116] dbplyr_2.1.1
#> [117] MASS_7.3-54
#> [118] rappdirs_0.3.3
#> [119] boot_1.3-28
#> [120] Matrix_1.3-4
```

```
#> [121] readr_2.0.2
#> [122] Gviz_1.36.2
#> [123] igraph_1.2.6
#> [124] TxDb.Hsapiens.UCSC.hg19.knownGene_3.2.2
#> [125] pkgconfig_2.0.3
#> [126] GenomicAlignments_1.28.0
#> [127] coin_1.4-1
#> [128] foreign_0.8-81
#> [129] xml2_1.3.2
#> [130] ggtree_3.0.4
#> [131] yulab.utils_0.0.2
#> [132] stringr_1.4.0
#> [133] VariantAnnotation_1.38.0
#> [134] digest_0.6.28
#> [135] strucchange_1.5-2
#> [136] rmarkdown_2.11
#> [137] fastmatch_1.1-3
#> [138] tidytree_0.3.5
#> [139] htmlTable_2.2.1
#> [140] annotatr_1.18.1
#> [141] restfulr_0.0.13
#> [142] curl_4.3.2
#> [143] gtools_3.9.2
#> [144] shiny_1.7.1
#> [145] Rsamtools_2.8.0
#> [146] modeltools_0.2-23
#> [147] rjson_0.2.20
#> [148] jsonlite_1.7.2
#> [149] lifecycle_1.0.1
#> [150] nlme_3.1-153
#> [151] viridisLite_0.4.0
#> [152] fansi_0.5.0
#> [153] pillar_1.6.3
#> [154] lattice_0.20-45
#> [155] plotrix_3.8-2
#> [156] KEGGREST_1.32.0
#> [157] fastmap_1.1.0
#> [158] httr_1.4.2
#> [159] survival_3.2-13
#> [160] GO.db_3.13.0
#> [161] interactiveDisplayBase_1.30.0
#> [162] glue_1.4.2
#> [163] png_0.1-7
#> [164] BiocVersion_3.13.1
#> [165] bit_4.0.4
#> [166] ggforce_0.3.3
#> [167] stringi_1.7.5
#> [168] blob_1.2.2
#> [169] AnnotationHub_3.0.1
#> [170] caTools_1.18.2
#> [171] latticeExtra_0.6-29
#> [172] memoise_2.0.0
#> [173] dplyr_1.0.7
```

```
#> [174] ape_5.5
```
