## Supplementary material for "epiRomics: a multi-omics R package to identify and visualize enhancers": EpiRomics Manual

### Package ‘epiRomics’

September 30, 2021

**Type** Package

**Title** Epigenomic Analysis Package Built for R (epiRomics)

**Version** 0.1.3

**Maintainer** Alex M. Mawla <>

**Description** A package designed to integrate various levels of epigenomic information, including, but not limited to ChIP and Histone next-generation sequencing. Regulatory network analysis can be done by using combinatory approaches to infer regions of significance, such as enhancers. Downstream analysis can identify co-occurance of epigenomic data located at regions of interest. Finally, this package allows for various results to be visualized. This package is currently in development. Please contact <> for suggestions, feedback, or bug reporting.

**License** MIT + file LICENSE

**Depends** R (>= 4.0.0)

**biocViews**

**Imports** AnnotationDbi (>= 1.54.1),
annotatr (>= 1.18.1),
BiocGenerics (>= 0.38.0),
BiocManager (>= 1.30.16),
ChIPseeker (>= 1.28.3),
data.table (>= 1.12.8),
GenomeInfoDb (>= 1.28.1),
GenomicFeatures (>= 1.44.1),
GenomicRanges (>= 1.44.0),
Gviz (>= 1.36.2),
igraph (>= 1.2.6),
IRanges (>= 2.26.0),
methods (>= 4.0.2),
party (>= 1.3.3),
plyr (>= 1.8.6),
rtracklayer (>= 1.52.1),
utils

**Suggests** covr,
org.Hs.eg.db (>= 3.13.0),
TxDb.Hsapiens.UCSC.hg38.knownGene (>= 3.13.0),
knitr,

testthat (>= 3.0.0),
rmarkdown,
enrichplot,
pkgdown

**VignetteBuilder** knitr

**Remotes** GuangchuangYu/enrichplot,
GuangchuangYu/ChipSeeker

**Config/testthat/edition** 3

**ByteCompile** true

**Encoding** UTF-8

**LazyData** true

**RoxygenNote** 7.1.2

### R topics documented:

**Index** **11**

---

`epiRomicsS4-class` *An S4 class to manage epiRomics databases and downstream results*

---

##### Description

An S4 class to manage epiRomics databases and downstream results

##### Slots

`annotations` GRanges

`meta` data.frame

`txdb` txdb string name

`organism` org.db string name

`genome` genome name, e.g. 'mm10' or 'hg38'

#### epiRomics_build_dB *Build epiRomics database*

##### Description

Build epiRomics database

##### Usage

```
epiRomics_build_dB(
  epiRomics_db_file,
  txdb_organism,
  epiRomics_genome,
  epiRomics_organism
)
```

##### Arguments

`epiRomics_db_file`
: character string of path to properly formatted csv file containing epigenetic data. [See vignette for more details]

`txdb_organism`
: a character string containing the TxDB associated with your data.

`epiRomics_genome`
: a character string containing the genome associated with your data. e.g. "mm10" or "hg19".

`epiRomics_organism`
: a character string containing the org.db associated with your data.

##### Value

Variable of class epiRomics for further downstream analysis

#### epiRomics_build_dB_2 *Build epiRomics database*

##### Description

Build epiRomics database

##### Usage

```
epiRomics_build_dB_2(
  epiRomics_db_file,
  txdb_organism,
  epiRomics_genome,
  epiRomics_organism
)
```

#### Arguments

epiRomics_db_file
: character string of path to properly formatted csv file containing epigenetic data. [See vignette for more details]

txdb_organism
: a character string containing the TxDB associated with your data.

epiRomics_genome
: a character string containing the genome associated with your data. e.g. "mm10" or "hg38".

epiRomics_organism
: a character string containing the org.db associated with your data.

#### Value

Variable of class epiRomics for further downstream analysis

---

### epiRomics_chromatin_tracks

*Visualizes chromatin availability*

---

#### Description

Visualizes chromatin availability

#### Usage

```
epiRomics_chromatin_tracks(
  epiRomics_gene_name,
  epiRomics_dB,
  epiRomics_track_connection
)
```

#### Arguments

epiRomics_gene_name
: character of name of gene to visualize

epiRomics_dB
: epiRomics class database containing all data initially loaded

epiRomics_track_connection
: data frame containing bigwig track locations and their names

#### Value

GViz plot

---

`epiRomics_enhanceosome` *Identifies putative enhanceosome regions by cross-referencing candidate enhancer regions against co-TF enrichment*

---

#### Description

Identifies putative enhanceosome regions by cross-referencing candidate enhancer regions against co-TF enrichment

#### Usage

```
epiRomics_enhanceosome(epiRomics_putative_enhancers, epiRomics_dB)
```

#### Arguments

`epiRomics_putative_enhancers`
epiRomics class database containing putative enhancer calls

`epiRomics_dB` epiRomics class database containing all data initially loaded

#### Value

Variable of class epiRomics further exploring candidate enhanceosome regions using co-ChIP hits

---

`epiRomics_enhancers` *Identifies putative enhancer regions utilizing select histone marks*

---

#### Description

Identifies putative enhancer regions utilizing select histone marks

#### Usage

```
epiRomics_enhancers(
  epiRomics_dB,
  epiRomics_histone_mark_1 = "h3k4me1",
  epiRomics_histone_mark_2 = "h3k27ac"
)
```

#### Arguments

`epiRomics_dB` epiRomics class database containing all data initially loaded

`epiRomics_histone_mark_1`
name of first histone mark, must match name in epiROmics_dB@meta, default set to h3k4me1

`epiRomics_histone_mark_2`
name of second histone mark, must match name in epiROmics_dB@meta default set to h3k27ac

#### Value

Variable of class epiRomics further exploring candidate enhancer regions identified after histone integration

---

`epiRomics_enhancers_filter`
*Filters putative enhancers called by epiRomics_enhancers by crossing against curated FANTOM data*

---

#### Description

Filters putative enhancers called by epiRomics_enhancers by crossing against curated FANTOM data

#### Usage

```
epiRomics_enhancers_filter(
  epiRomics_putative_enhancers,
  epiRomics_dB,
  epiRomics_type = "mm10_custom_fantom"
)
```

#### Arguments

| | |
|---|---|
| `epiRomics_putative_enhancers` | epiRomics class database containing putative enhancer calls |
| `epiRomics_dB` | epiRomics class database containing all data initially loaded |
| `epiRomics_type` | epiRomics reference containing database to validate putative enhancers against |

#### Value

Variable of class epiRomics with filtered candidate enhancer regions

---

`epiRomics_enhancer_predictor_test`
*Interrogates various histone marks against a curated database to determine which are most informative*

---

#### Description

Interrogates various histone marks against a curated database to determine which are most informative

#### Usage

```
epiRomics_enhancer_predictor_test(
  epiRomics_dB,
  epiRomics_histone = "h3k4me1",
  epiRomics_curated_database = "fantom"
)
```

##### Arguments

`epiRomics_dB` epiRomics class database containing all data initially loaded

`epiRomics_histone`
name or vector of histone mark(s), must match name in epiROmics_dB@meta, default set to h3k4me1

`epiRomics_curated_database`
database to test histone marks against, must match name in epiROmics_dB@meta default set to fantom

##### Value

Variable of class dataframe further exploring top histone marks that may determine enhancer regions

---

#### `epiRomics_predictors` *Predicts TF behavior in association with enhanceosome presence in accordance with functional annotation*

---

##### Description

Predicts TF behavior in association with enhanceosome presence in accordance with functional annotation

##### Usage

```
epiRomics_predictors(epiRomics_putative_enhanceosome)
```

##### Arguments

`epiRomics_putative_enhanceosome`
epiRomics class database containing putative enhanceosome calls

##### Value

Returned decision tree available for plotting

---

#### `epiRomics_regions_of_interest` *Evaluates whether regions of interest derived from external experiments, such as ATAC-Seq, correspond with enhanceosome regions*

---

##### Description

Evaluates whether regions of interest derived from external experiments, such as ATAC-Seq, correspond with enhanceosome regions

##### Usage

```
epiRomics_regions_of_interest(
  epiRomics_putative_enhanceosome,
  epiRomics_test_regions
)
```

#### Arguments

`epiRomics_putative_enhanceosome`
: epiRomics class database containing putative enhanceosome calls

`epiRomics_test_regions`
: GRanges containing regions of interest

#### Value

Variable of class epiRomics with enhanceosome regions overlapping with regions of interest

---

epiRomics_region_tracks *Visualizes chromatin availability at custom regions*

---

#### Description

Visualizes chromatin availability at custom regions

#### Usage

```
epiRomics_region_tracks(
  epiRomics_region,
  epiRomics_dB,
  epiRomics_track_connection
)
```

#### Arguments

`epiRomics_region`
: GRanges of region to visualize

`epiRomics_dB` epiRomics class database containing all data initially loaded

`epiRomics_track_connection`
: data frame containing bigwig track locations and their names

#### Value

GViz plot

---

`epiRomics_track_layer` *Visualizes data from epiRomics results*

---

#### Description

Visualizes data from epiRomics results

#### Usage

```
epiRomics_track_layer(
  epiRomics_putative_enhanceosome,
  epiRomics_index,
  epiRomics_dB,
  epiRomics_track_connection,
  epiRomics_keep_epitracks = TRUE
)
```

#### Arguments

`epiRomics_putative_enhanceosome`
: epiRomics class database containing putative enhanceosome calls

`epiRomics_index`
: numeric of row value from epiRomics_putative_enhanceosome to visualize

`epiRomics_dB`
: epiRomics class database containing all data initially loaded

`epiRomics_track_connection`
: data frame containing bigwig track locations and their names

`epiRomics_keep_epitracks`
: logical indicating whether to show enhancer and chip tracks, default is TRUE

#### Value

GViz plot

---

`epiRomics_track_layer_human` *Visualizes data from epiRomics results*

---

#### Description

Visualizes data from epiRomics results

#### Usage

```
epiRomics_track_layer_human(
  epiRomics_putative_enhanceosome,
  epiRomics_index,
  epiRomics_dB,
  epiRomics_track_connection,
  epiRomics_keep_epitracks = TRUE
)
```

#### Arguments

`epiRomics_putative_enhanceosome`
: epiRomics class database containing putative enhanceosome calls

`epiRomics_index`
: numeric of row value from epiRomics_putative_enhanceosome to visualize

`epiRomics_dB`
: epiRomics class database containing all data initially loaded

`epiRomics_track_connection`
: data frame containing bigwig track locations and their names

`epiRomics_keep_epitracks`
: logical indicating whether to show enhancer and chip tracks, default is TRUE

#### Value

GViz plot

### Index

epiRomics_build_dB, 3
epiRomics_build_dB_2, 3
epiRomics_chromatin_tracks, 4
epiRomics_enhanceosome, 5
epiRomics_enhancer_predictor_test, 6
epiRomics_enhancers, 5
epiRomics_enhancers_filter, 6
epiRomics_predictors, 7
epiRomics_region_tracks, 8
epiRomics_regions_of_interest, 7
epiRomics_track_layer, 9
epiRomics_track_layer_human, 9
epiRomicsS4 *(*epiRomicsS4-class*)*, 2
epiRomicsS4-class, 2
